# (p)ppGpp directly regulates translation initiation during entry into quiescence

**DOI:** 10.1101/807917

**Authors:** Simon Diez, Jaewook Ryu, Kelvin Caban, Ruben L. Gonzalez, Jonathan Dworkin

## Abstract

Many bacteria exist in a state of metabolic quiescence where they must minimize energy consumption so as to maximize available resources over a potentially extended period of time. As protein synthesis is the most energy intensive metabolic process in a bacterial cell, it would be an appropriate target for downregulation during the transition from growth to quiescence. We find that when *Bacillus subtilis* exits growth, a subpopulation of cells emerges with very low levels of protein synthesis dependent on synthesis of the nucleotides (p)ppGpp. We show that (p)ppGpp inhibits protein synthesis *in vivo* and *in vitro* by preventing the allosteric activation of the essential GTPase Initiation Factor 2 (IF2) during translation initiation. Finally, we demonstrate that IF2 is an authentic *in vivo* target of (p)ppGpp during the entry into quiescence, thus providing a mechanistic basis for the observed attenuation of protein synthesis.

## Introduction

Most microbial life exists in a non-proliferating state of quiescence that enables survival during nutrient limitation and in stressful environments (Lennon and Jones, 2011; Rittershaus et al., 2013). A major challenge facing a quiescent cell is how to minimize energy consumption so as to maximize available resources over a potentially extended period of time. As protein synthesis is the most energy intensive metabolic process in a cell, accounting for as much as ∼70% of total energy consumption in bacteria (Tempest and Neijssel, 1984), it would be an appropriate target for downregulation during the transition from growth to quiescence. Metabolic conditions that may be present during this transition such as amino acid limitation would reduce protein synthesis, but whether it is also actively inhibited remains an open question.

Several highly conserved, essential GTPases participate in translation (Maracci and Rodnina, 2016). For example, IF2 is involved in initiation and EF-Tu and EF-G are involved in elongation. Thus, as key factors in translation, these proteins are appealing regulatory targets for quiescence-dependent attenuation of protein synthesis. In the bacterium *B. subtilis* undergoing sporulation, a well-characterized developmental pathway leading to quiescence, phosphorylation of EF-Tu inhibits its ability to hydrolyze GTP, an activity essential for its function in translation, and thereby blocks protein synthesis (Pereira et al., 2015). How else could these GTPases be targeted? Binding of a ligand other than GTP to the GTP binding site could affect protein function. For example, *in vitro*, the hyperphosphorylated nucleotides guanosine tetraphosphate and penta- phosphate, derived from GDP and GTP, respectively, can bind and thereby inhibit translational GTPases (Hamel and Cashel, 1974). These so-called alarmones, collectively termed (p)ppGpp, are the major mediators of the stringent response, the mechanism used by bacteria to coordinate a response to cell stresses including nutrient deprivation (Gaca et al., 2015; Steinchen and Bange, 2016). For example, (p)ppGpp synthesis occurs during the entry into stationary phase and in the presence of amino acid limitation (Steinchen and Bange, 2016). (p)ppGpp is responsible for the down-regulation of metabolic processes including transcription, replication, and GTP synthesis that are mediated by a direct interaction with RNA polymerase (Artsimovitch et al., 2004; Ross et al., 2016), DNA primase (Wang et al., 2007) and the GTP biosynthetic enzymes HprT and Gmk (Kriel et al., 2012), respectively.

Overexpression of a truncated RelA protein that synthesizes (p)ppGpp in the absence of amino acid limitation results in a rapid decrease in ^35^S-methionine incorporation (Svitil et al., 1993), consistent with an inhibitory effect of (p)ppGpp on translation. However, further investigation of a direct effect of (p)ppGpp on translation has been complicated by several factors. First, (p)ppGpp affects synthesis of ribosomal proteins in *E. coli* (Lindahl et al., 1976). Whether this effect is direct has been difficult to assess given the well-established effect of (p)ppGpp on transcription by *E. coli* polymerase. Second, studies examining the effect of (p)ppGpp produced by RelA use conditions that produce uncharged tRNAs in order to stimulate RelA (Arenz et al., 2016; Brown et al., 2016; Haseltine and Block, 1973; Loveland et al., 2016; O’Farrell, 1978). Since uncharged tRNAs directly arrest translation, it has been difficult to differentiate this effect from a direct effect of (p)ppGpp on translation.

*In vitro*, (p)ppGpp inhibits translation in a manner similar to that observed with non-hydrolyzable GTP analogs (Wagner and Kurland, 1980), suggesting that (p)ppGpp is targeting a translational GTPase. Consistently, (p)ppGpp inhibits the GTPase activity of IF2 and EF-Tu (Hamel and Cashel, 1974) as well as of GTPases that mediate other aspects of translation such as ribosome assembly (Corrigan et al., 2016; Pausch et al., 2018), by acting as competitive inhibitors. (p)ppGpp is capable of binding translational GTPases including EF-Tu, EF-G (Rojas et al., 1984), IF2 (Milon et al., 2006; Mitkevich et al., 2010) and the ribosome assembly GTPase ObgE (Persky et al., 2009) at affinities that are commensurate with the *in vivo* levels of (p)ppGpp observed following stringent response induction, consistent with the enzymes being *in vivo* targets. Of note, the affinity of IF2 for (p)ppGpp as compared to EF-G (Mitkevich et al., 2010) and the observation that (p)ppGpp interferes with IF2 function (Milon et al., 2006) suggests that IF2 may be a principal target *in vivo*. However, this is not known.

Here we show that during the transition into a non-proliferative state, *Bacillus subtilis* exhibits a large reduction in protein synthesis that is dependent on (p)ppGpp. We further show that both *in vivo* and *in vitro*, (p)ppGpp inhibits protein synthesis by targeting translation. Next, we identify mutations in IF2 which allow us to demonstrate *in vitro* that it is a direct target of (p)ppGpp during translation. We then show that binding of ppGpp fails to allosterically stabilize a conformation of IF2 that is typically triggered by binding of GTP and that enables IF2 to stably bind the ribosomal small, or 30S, subunit initiation complex (IC) and catalyze rapid joining of the ribosomal large, or 50S, subunit to the 30S IC. Finally, we demonstrate *in vivo* that binding of (p)ppGpp to IF2 mediates the observed (p)ppGpp-dependent inhibition of protein synthesis.

## Results

### Protein synthesis is inhibited in a (p)ppGpp dependent manner during stationary phase

*B. subtilis* grows exponentially in LB until a stereotypic cell density, presumably dictated by nutrient availability. After this point, growth occurs more slowly (non-exponentially) in the transition phase which culminates in the non-proliferative state of stationary phase (Figure 1A). We assayed protein synthesis during different growth phases by measuring incorporation of the puromycin analog *O*-propargyl-puromycin (OPP) that can be visualized and quantified in single cells following addition of a fluorophore using click chemistry (Liu et al., 2012). Incorporation of OPP results in the accumulation of fluorescently tagged nascent polypeptide chains that directly reflects the rate of translation (Liu et al., 2012). Although puromycin causes premature termination of protein synthesis resulting from its incorporation into the nascent polypeptide chain, global protein synthesis occurs even following addition of OPP under our conditions (Figure S1). Other methods for measuring protein synthesis exist, but they rely on growing cells in the absence of amino acids (specifically methionine). This poses a particular problem for strains which exhibit autotrophies to these amino acids, such as those lacking (p)ppGpp (Kriel et al., 2014). Thus, the use of OPP allows us to examine protein synthesis under conditions where amino acids are not specifically limiting as well as at the level of single cells.

**Figure 1.**
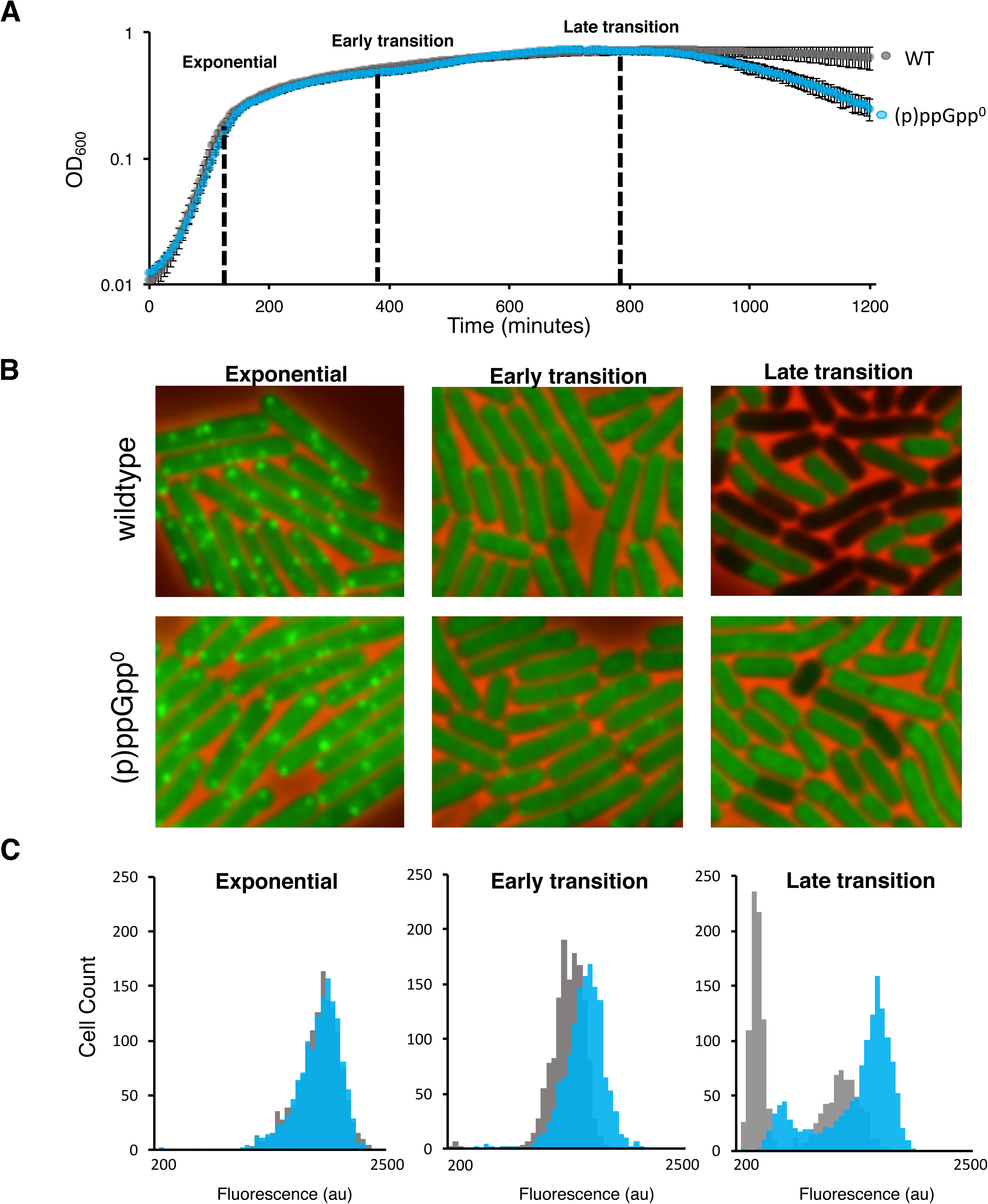
(p)ppGpp mediates inhibition of protein synthesis upon departure from exponential growth. *B. subtilis* (JDB 1772) cells were labeled with OPP at different time points following exponential phase. **(A)** *B. subtilis* wildtype (WT) and isogenic (p)ppGpp^0^ (JDB 4294) strains grow equivalently to each other during exponential and transition phases but (p)ppGpp^0^ strain lyses upon entry into stationary phase (means ± SDs). **(B)** Representative pictures of WT and (p)ppGpp^0^ strains labelled with OPP at different time points. **(C)** Distributions of mean-cell fluorescence of WT and (p)ppGpp^0^ cells. Time points in (B) and (C) are indicated by black dashed lines in (A). See also Figure S1 and Figure S2

We labeled *B. subtilis* cultures with OPP at a series of time points (Figure 1A; dashed black lines). As expected, cells exhibited a progressive decrease in OPP incorporation soon after departure from exponential growth (early transition) that continued as the cells grew slowly in late transition (Figure 1B, top). This trend is apparent in the average cellular fluorescence at these time points as well as in stationary phase (Figure S2A, gray bars). However, at the late transition time point, a substantial fraction of cells in the population appeared to lose all fluorescent signal (Figure 1B, top). This loss of signal indicates an absence of protein synthesis, resulting in a population whose distribution of the rates of protein synthesis is roughly bimodal at the single cell level (Figure 1C; gray). This heterogeneity is consistent with the inhibition of protein synthesis being a direct and acute effect in a sub-population of cells rather than the consequence of amino acid limitation in the growth medium which would be expected to cause a homogenous decrease across the entire population. That is, a sub-population of cells in the late transition phase culture exhibited a near total inhibition of protein synthesis whereas a separate sub-population maintained a noticeably higher level of protein synthesis (Figure 1C, gray).

We speculated that a mechanism known to control growth rate might be important for the shutdown of protein synthesis in post-exponential growth. An attractive candidate is (p)ppGpp, a molecule involved in regulating diverse processes that affect cells growing under sub-optimal conditions (Potrykus et al., 2011). Furthermore, many bacteria synthesize (p)ppGpp when they depart from exponential phase (Boes et al., 2008; Murray et al., 2003). To investigate the possible role of (p)ppGpp, we utilized a strain that lacks (p)ppGpp *via* genetic deletion of the three known *B. subtilis* (p)ppGpp synthetases, *relA*, *sasA*, and *sasB* (Nanamiya et al., 2008; Srivatsan et al., 2008), which we will refer to as (p)ppGpp^0^. Under our growth conditions, the (p)ppGpp^0^ strain grows equivalently to the parent wildtype strain during exponential phase as well as early and late transition phase. As above, we assayed protein synthesis of the (p)ppGpp^0^ strain by measuring OPP incorporation. Exponential phase OPP incorporation of this strain is indistinguishable from the parent (Figure 1B, C). In contrast, early in transition phase, some wildtype cells incorporate substantially less OPP than (p)ppGpp^0^ cells. This trend continues and at a time point late in transition phase there are significantly less (p)ppGpp^0^ cells (blue) which have decreased their protein synthesis compared to the wildtype parent (gray) (Figure 1B, C; Figure S2A, B). Thus, (p)ppGpp is necessary for the observed inhibition of protein synthesis during late transition phase even though cells lacking (p)ppGpp do not display a different growth rate up to this point.

### (p)ppGpp is sufficient to inhibit protein synthesis

Since (p)ppGpp^0^ cells failed to decrease protein synthesis during the transition phase, we investigated whether production of (p)ppGpp is sufficient to inhibit protein synthesis. The *B. subtilis* (p)ppGpp synthetase *sasA* was placed under the control of a xylose-inducible promoter in a strain lacking RelA, the bifunctional (p)ppGpp synthetase and hydrolase, as well the accessory (p)ppGpp synthetases SasA and SasB. As previously observed (Tagami et al., 2012), induction of *sasA* in the presence of these mutations results in a decrease in growth rate which culminates in cessation of growth (Figure 2A). To determine how (p)ppGpp affects protein synthesis, cells were labeled with OPP at 30-minute time intervals following xylose addition. At the time of addition (T_0_), there is not a significant difference in protein synthesis between the induced and un-induced cultures. However, at 30 minutes following inducer addition (T_30_) and at later times (T_60_, T_90_), protein synthesis is significantly reduced in the induced cultures as compared to the un-induced cultures (Figure 2B, C).

**Figure 2.**
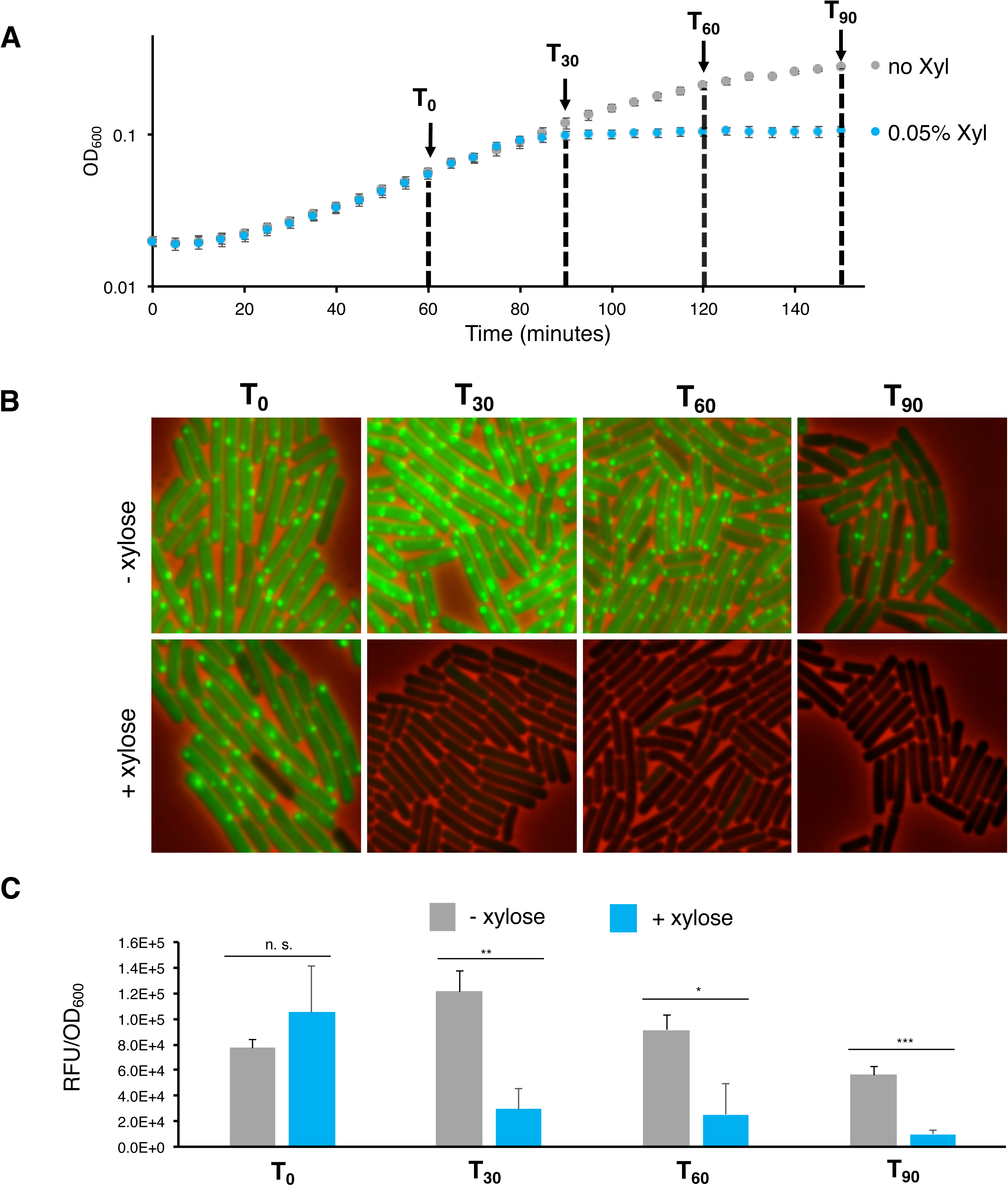
(p)ppGpp is sufficient to inhibit growth and protein synthesis. SasA was expressed during exponential phase growth and cells were labelled with OPP. **(A)** A P*_xyl_-sasA* strain (JDB 4295) was grown in duplicate and 0.05% xylose was added after 60 min of growth (T_0_) and growth was monitored for 90 min post induction (means ± SDs). **(B)** Representative pictures of OPP-labelled induced and un-induced cultures of P*_xyl_-sasA* strain at time points post induction. **(C)** Total fluorescence of OPP labelled induced and un-induced cultures of P*_xyl_-sasA* strain at different time points post induction. Time points in (B) and (C) are designated by black dashed lines in (A) (means ± SDs). n.s. p > 0.05, *p < 0.05, **p < 0.01, ***p < 0.001

### (p)ppGpp inhibits translation *in vivo*

We reasoned that measuring both the synthesis of a single protein and the transcription of its gene would allow us to definitively demonstrate that inhibition of protein synthesis was due to inhibition of translation not transcription. We chose the *B. subtilis* P*_veg_* promoter which is insensitive to (p)ppGpp (Krasny and Gourse, 2004) and firefly luciferase as a reporter protein because its half-life in *B. subtilis* is only 6 minutes (Mirouze et al., 2011), so measurements of its activity closely reflect the kinetics of its synthesis. We grew strains that contained an inducible copy of *sasA* (P*_xyl_*-*sasA*) as well the P*_veg_*-*luc* reporter and compared luciferase activity in the presence and absence of inducer. While luciferase activity is easily detected during exponential growth, *sasA* induction is quickly followed by a drastic decrease in the luciferase activity per cell as compared to cells in the absence of induction (Figure 3A). This decrease in luciferase activity does not occur at the level of transcription as the level of *luc* mRNA on a per cell level is similar in the induced and un-induced cultures (Figure 3B). Furthermore, this decrease is not dependent on changes in levels of 16S ribosomal RNA (rRNA) which would affect ribosome abundance (Figure 3B). Thus, (p)ppGpp is sufficient to decrease synthesis of a particular protein without affecting transcription of its mRNA, consistent with a direct effect on translation.

**Figure 3.**
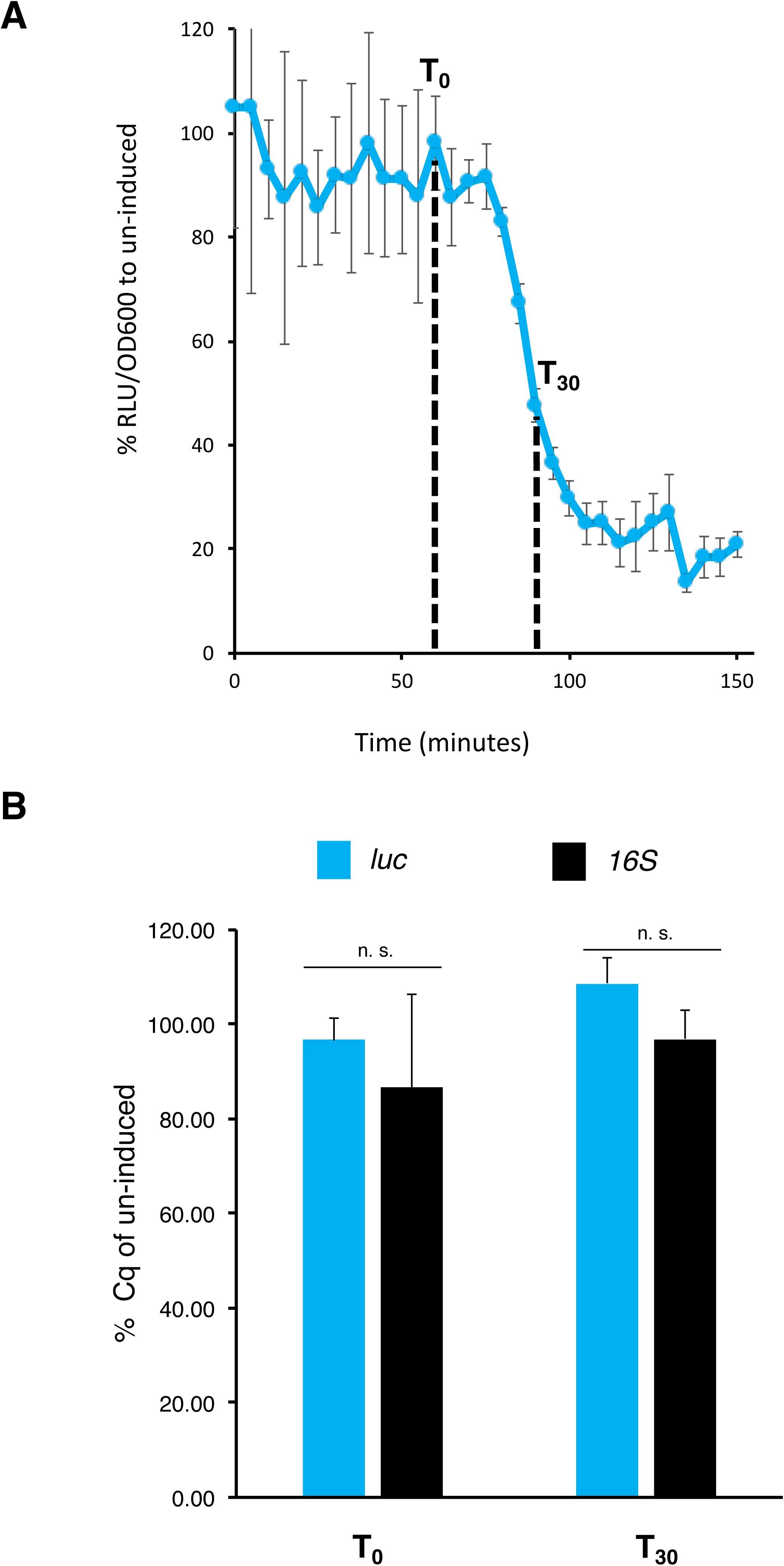
(p)ppGpp directly inhibits translation *in vivo*. Luminescence produced by a luciferase reporter protein was measured during exponential phase growth before and after *sasA* expression. **(A)** *P_xyl_-sasA P_veg_-luc* strain (JDB4296) was grown in duplicate and 0.05% xylose was added after 60 min of growth (T_0_) and growth and luminescence were measured for 90 min post induction of *sasA* (means ± SDs). **(B)** Direct inhibition of translation of the luciferase reporter was verified by measuring *luc* mRNA and *16S* rRNA levels both at the time of induction and a time point when the *luc* reporter was significantly inhibited (T_0_ and T_30_ respectively) using RT-qPCR (means ± SDs). n.s. p > 0.05, *p < 0.05, **p < 0.01, ***p < 0.001

### (p)ppGpp inhibits translation *in vitro*

We extended these *in vivo* observations to *in vitro* experiments using the PURExpress *in vitro* reconstituted, coupled transcription-translation system (NEB) which utilizes a defined mix of purified transcription and translation components to transcribe and translate a specific mRNA (Shimizu et al., 2014). Addition of ppGpp to PURExpress reactions inhibited synthesis of a reporter protein (CotE) in a dose-dependent fashion (Figure 4). Concentrations of ppGpp sufficient to significantly inhibit CotE synthesis (∼1 mM) are similar to levels of (p)ppGpp observed during stringent response induction in *E. coli* (Buckstein et al., 2008). qRT-PCR analysis demonstrated that ppGpp had no effect on *cotE* transcription at levels where CotE synthesis was significantly impaired (Figure S3), consistent with the known insensitivity of the T7 RNA polymerase used in the PURExpress reaction to ppGpp (Friesen and Fiil, 1973). Thus, similar to the *in vivo* data (Figure 3), these *in vitro* data demonstrate that ppGpp directly inhibits translation.

**Figure 4.**
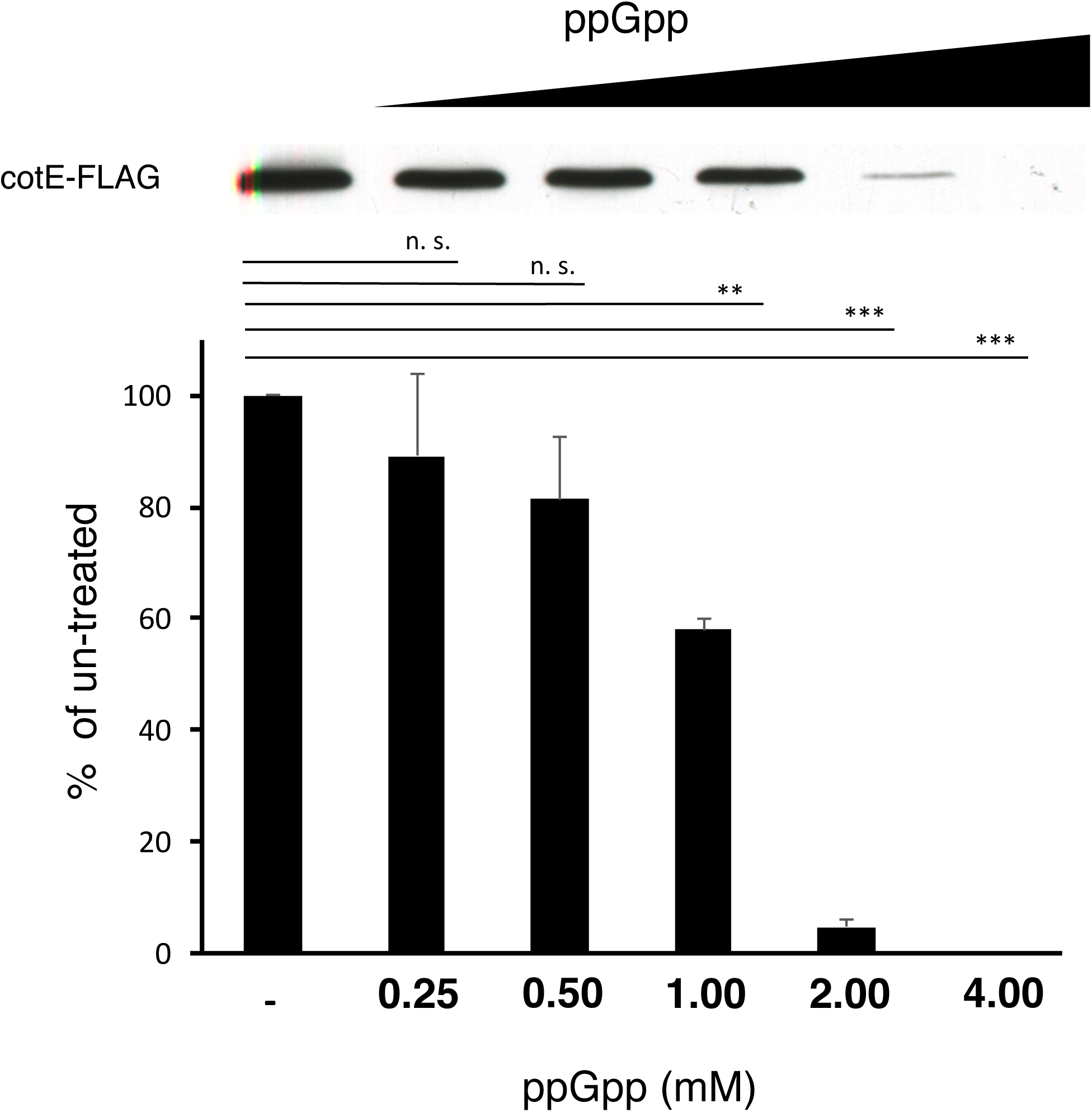
ppGpp directly inhibits translation *in vitro*. Protein synthesis in the presence of increasing concentrations of ppGpp was measured using the PURExpress *in vitro* reconstituted, coupled transcription-translation system (NEB). Production of CotE-FLAG was measured via Western blot with α-FLAG (means ± SDs). n.s. p > 0.05, *p < 0.05, **p < 0.01, ***p < 0.001 See also Figure S3

### IF2 is a target of (p)ppGpp

Given these observations, we wished to identify the component(s) of the translation machinery that is (are) targeted by ppGpp. The translational GTPases EF-Tu, EF-G and IF2 as well as the ribosome associated GTPases including Obg (Buglino et al., 2002; Feng et al., 2014), RsgA (Corrigan et al., 2016; Zhang et al., 2018), RbgA (Corrigan et al., 2016; Pausch et al., 2018), Era (Corrigan et al., 2016), and HflX (Corrigan et al., 2016; Zhang et al., 2018) have all been reported to bind (p)ppGpp. However, (p)ppGpp inhibits protein synthesis by the PURExpress system (Figure 4), which contains only IF2, EF-Tu, and EF-G, so inhibition of one or more of these proteins is likely sufficient to account for the *in vivo* inhibitory effect of (p)ppGpp on translation.

Initiation is a key point of translational control during nutrient limitation in yeast cells (Hinnebusch, 2005) and since ppGpp inhibits IF2 *in vitro* (Legault et al., 1972; Milon et al., 2006), we first investigated whether IF2 was a target of (p)ppGpp under our conditions. To do this, we attempted to identify mutations in IF2 that would affect (p)ppGpp binding without disrupting GTP binding sufficiently to impair normal function. This goal was inspired by the demonstration that replacing the *B. subtilis* GTP synthesis enzyme GMK that is sensitive to (p)ppGpp with the *E. coli* homolog that is insensitive to (p)ppGpp altered the *B. subtilis* physiological response to elevated (p)ppGpp levels (Liu et al., 2015). However, the G1-G3 motifs of the G domain that interact with GTP are nearly identical between *E. coli* and *B. subtilis* IF2 (Verstraeten et al., 2011), so a total allele substitution strategy seemed unlikely to be similarly informative for IF2. Alternatively, *E. coli* EF-G and IF2 have differential affinity for (p)ppGpp (Mitkevich et al., 2010) and similar but not as extensively conserved G domains such that, if we could identify residue(s) that affect this difference, this information might allow us to construct a *B. subtilis* IF2 allele less sensitive to (p)ppGpp.

We focused on those IF2 residues which display a shift in NMR spectra upon binding of ppGpp as compared to GDP (Figure 5B, blue residues) (Milon et al., 2006) since GDP interacts with multiple residues in the G domain of IF2 (Wienk et al., 2012). We aligned the region containing those residues (*i.e.*, the G1 motif) with the homologous region in EF-G, which has lower affinity for (p)ppGpp than IF2 (Figure 5A). We noted that one of the blue residues in IF2, Gly-226, is an alanine residue (Ala-18) in EF-G (Figure 5B, red). In addition, the histidine residue (His-230) in IF2 that is adjacent to the two blue residues is an alanine (Ala-21) in EF-G (Figure 5B, red). These differences suggested that substituting the IF2 residues with the corresponding residues found in EF-G would affect the ability of IF2 to bind (p)ppGpp. We therefore compared the affinity of wild type and mutant IF2 for radiolabeled (p)ppGpp using the DRaCALA filter binding assay and observed that the double mutant IF2 (G226A H230A) bound (p)ppGpp significantly less well than the wildtype protein (Figure 5C).

**Figure 5.**
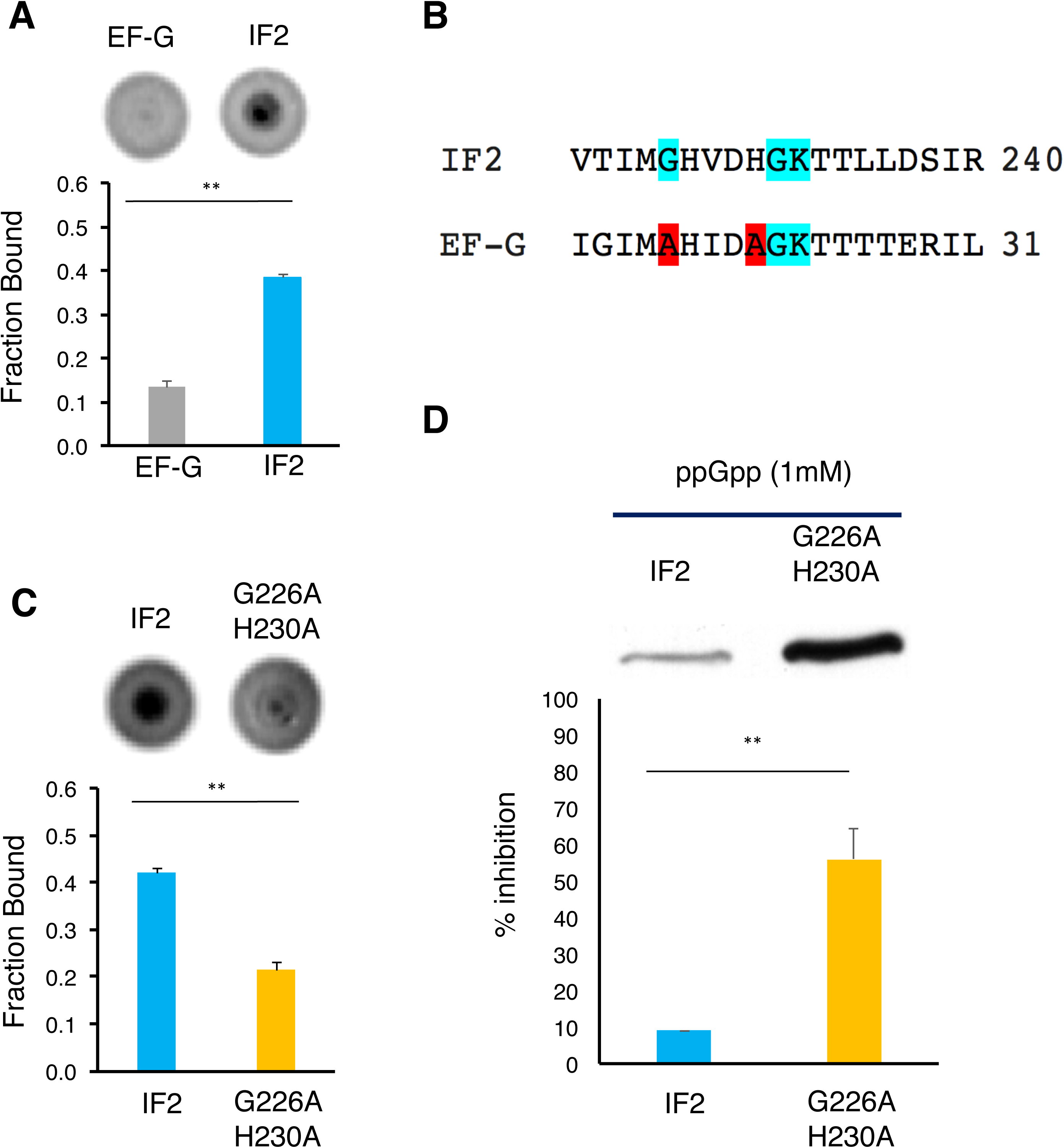
IF2 is a target of ppGpp. IF2 was validated *in vitro* as a direct target of ppGpp using IF2 mutations that reduce ppGpp binding. **(A)** Affinity of *B. subtilis* EF-G and IF2 for (p)ppGpp was compared using the differential radial capillary action of a ligand assay (DRaCALA) (Roelofs et al., 2011). (means ± SDs). **(B)** Alignment of G1 domains of *B. subtilis* IF2 and EF-G. Residues in blue denote those whose chemical shifts were previously identified to be most shifted upon binding of ppGpp *versus* GDP. Residues in red are those that were different in EF-G *versus* IF2 and that were used to engineer a mutant IF2 with reduced affinity for ppGpp (G226A H230A). **(C)** DRaCALA-based comparison of ppGpp affinity for WT and mutant IF2 (means ± SDs). **(D)** *in vitro* sensitivity of WT and mutant IF2 was assessed using the PURExpress *in vitro* reconstituted, coupled transcription-translation system (NEB). WT and mutant IF2 were added at equimolar amounts to separate PURExpress reactions in the presence of 1mM ppGpp and protein synthesis was monitored by Western blot (means ± SDs). n.s. p > 0.05, *p < 0.05, **p < 0.01, ***p < 0.001 See also Figure S4

To test the functional consequence of mutating these residues in *B. subtilis* IF2, we used a PURExpress kit that lacks IF2 (*Δ*IF2). We first confirmed that the *Δ*IF2 kit, which contains purified *E. coli* translation factors, works equivalently whether the added IF2 is derived from *E. coli* or *B. subtilis* (Figure S4B). When supplied as the sole source of IF2 in this reaction, the double mutant *B. subtilis* IF2 produced an equivalent amount of protein as wild type IF2 (Figure S4B) demonstrating that the slight reduction in GTP binding (Figure S4A) did not substantially affect IF2 function in translation. However, the double mutant *B. subtilis* IF2 was significantly less sensitive than its wild type counterpart to ppGpp (Figure 5D). Taken together, these results indicate that ppGpp binding to IF2 accounts for a substantial portion of the inhibition of translation by ppGpp.

### ppGpp fails to allosterically activate IF2 for rapid subunit joining

During 30S IC assembly, IF2 promotes binding of initiator tRNA (fMet-tRNA^fMet^) to the 30S subunit and uses its domain IV (dIV) to directly contact the *N*-formyl-methionine and aminoacyl acceptor stem of fMet-tRNA^fMet^, resulting in formation of an IF2-tRNA sub-complex on the intersubunit surface of the 30S IC (Simonetti et al., 2008). Subsequently, the presence of GTP in the G domain and recognition of fMet-tRNA^fMet^ by dIV ‘activate’ 30S IC-bound IF2 for rapid subunit joining (Pavlov et al., 2011). Using an IF2-tRNA single-molecule fluorescence resonance energy transfer (smFRET) signal that reports on the formation and conformational dynamics of the IF2-tRNA sub- complex (Wang et al., 2015), we have previously demonstrated that activation of IF2 for rapid subunit joining involves a GTP- and fMet-tRNA^fMet^-dependent conformational change of IF2 that results in an increase in the affinity of IF2 for the 30S IC and an increase in the rate of subunit joining (Caban et al., 2017). Notably, the presence of GTP in the G domain allosterically places dIV in close proximity to fMet-tRNA^fMet^, resulting in an IF2-tRNA sub-complex conformation characterized by a distribution of FRET efficiency (E_FRET_) values centered at a mean E_FRET_ value (<E_FRET_>) of ∼0.80 (Caban et al., 2017). In contrast, the presence of GDP in the G domain places dIV further from fMet-tRNA^fMet^, resulting in an IF2-tRNA sub-complex conformation characterized by an <E_FRET_> of ∼0.60, an ∼30-fold decrease in the affinity of IF2 for the 30S IC (Caban et al., 2017), and a ∼20-60 fold reduction in the rate of IF2-catalyzed subunit joining (Antoun et al., 2003) (Antoun et al., 2004; Pavlov et al., 2011).

Previously, Milón and colleagues demonstrated that ppGpp inhibits the ability of IF2 to catalyze 30S IC assembly and subunit joining, consequently inhibiting formation of the first peptide bond in the synthesis of a protein (Milon et al., 2006). To elucidate the structural basis through which (p)ppGpp inhibits these IF2 activities, we performed smFRET experiments using our IF2-tRNA smFRET signal to determine whether and how ppGpp influences the binding of *E. coli* IF2 to the *E. coli* 30S IC and the conformational dynamics of *E. coli* 30S IC-bound IF2 (Figure 6). We began by comparing the affinities of GTP-bound IF2 (IF2(GTP)) and ppGpp-bound IF2 (IF2(ppGpp)) for the 30S IC. As in our previous studies (Caban et al., 2017; Wang et al., 2015), E_FRET_ *versus* time trajectories recorded for individual 30S ICs were observed to fluctuate between a zero FRET state, corresponding to the IF2-free state of the 30S IC, and a non-zero FRET state, corresponding to the IF2-bound state of the 30S IC (Figure 6, third row). Initial inspection of these trajectories and the corresponding surface contour plots of the post-synchronized time evolution of population FRET reveals that, while IF2(GTP) exhibits relatively long-lived and stable binding events on the 30S IC, IF2(ppGpp) exhibits significantly shorter-lived and unstable binding events on the 30S IC (Figure 6, third and fourth rows). To quantitatively compare the affinities of IF2(GTP) and IF2(ppGpp) for the 30S IC, we extracted kinetic and thermodynamic parameters from the smFRET data describing the binding of IF2 to the 30S IC **(STAR Methods).** This analysis demonstrates that the equilibrium dissociation constant (*K*_d_) for IF2 binding to the 30S IC is ∼10- fold higher when IF2 is bound to ppGpp relative to GTP, demonstrating that IF2(ppGpp) has a significantly lower affinity for the 30S IC compared to IF2(GTP) (Table S1).

**Figure 6.**
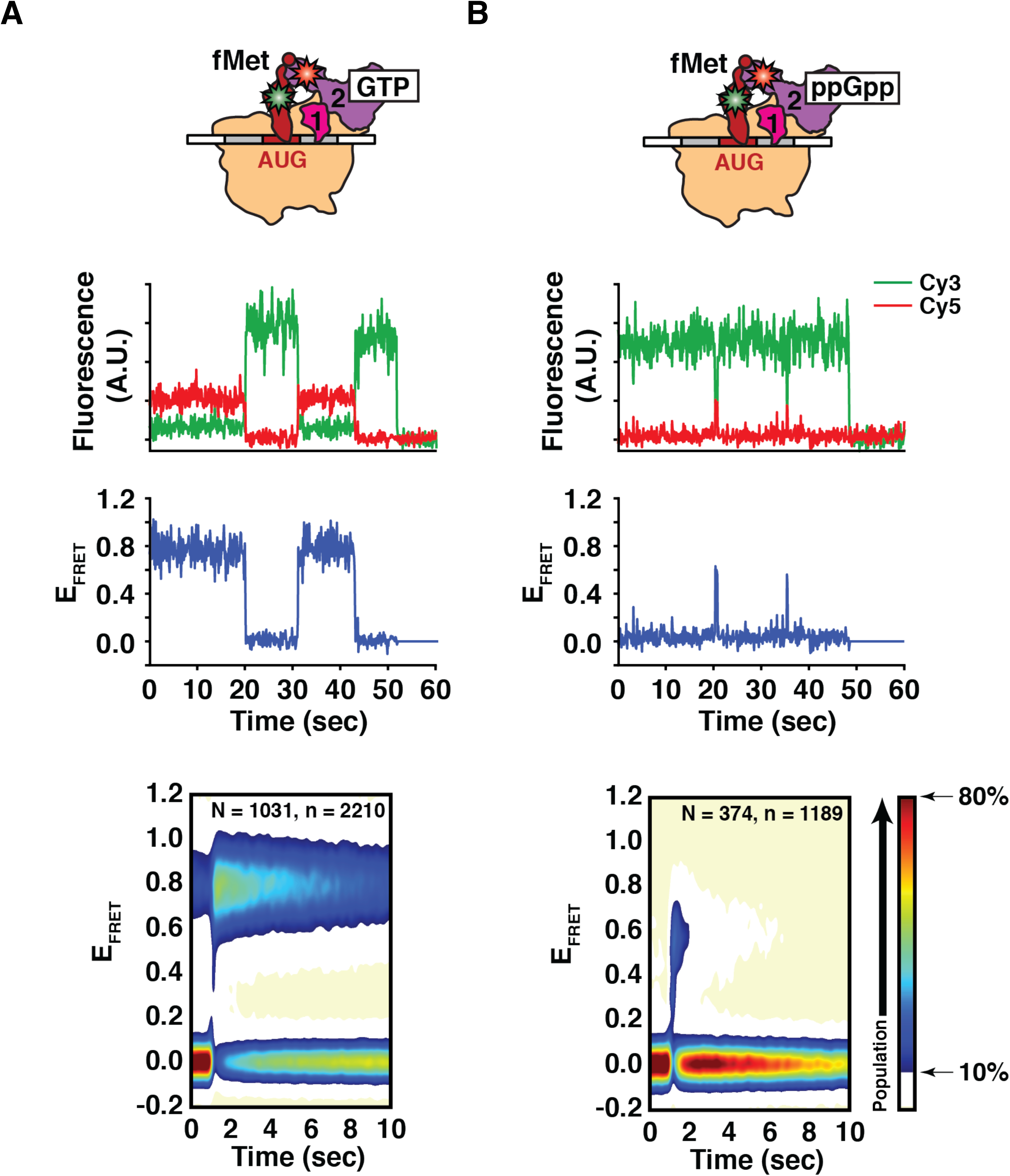
ppGpp *inhibits* IF2 function in catalyzing rapid 50S subunit joining. The binding of IF2 to the 30S IC and the conformation of 30S IC-bound IF2 in the presence of **(A)** GTP and **(B)** ppGpp were directly observed by single-molecule fluorescence resonance energy transfer (smFRET) using an IF2-tRNA smFRET signal. First row: Cartoon representations of 30S ICs assembled using Cy3 FRET donor fluorophore-labeled fMet-tRNA^fMet^ and Cy5 FRET acceptor fluorophore-labeled IF2(GTP) or IF2(ppGpp). Second row: Plots of Cy3 (green) and Cy5 (red) fluorescence emission intensity *versus* time trajectories. Third row: Plots of the E_FRET_ *versus* time trajectories corresponding to the plots of Cy3 and Cy5 fluorescence intensity trajectories in the second row. Fourth row: Surface contour plots of the post-synchronized time evolution of population FRET. These plots are generated by superimposing the E_FRET_ *versus* time trajectories of individual IF2 binding events such that the start of each event is computationally post- synchronized to time = 0 sec, thereby allowing visualization of the time evolution of population FRET for the entire population of IF2 binding events. “N” indicates the total number of individual 30S ICs analyzed and “n” indicates the total number of individual IF2 binding events analyzed. See also Figure S5 and Table S1

We then proceeded to compare the conformations of IF2(ppGpp) and IF2(GTP) on the 30S IC by plotting histograms of the E_FRET_ values observed under each condition (Figure S5). These histograms exhibit two distinct peaks centered at a zero <E_FRET_> and non-zero <E_FRET_> corresponding to the IF2-free and IF2-bound states of the 30S IC, respectively. Consistent with our previous studies, we observed a single non-zero peak centered at an <E_FRET_> of ∼0.80 for 30S IC-bound IF2(GTP), corresponding to a distance between our labeling positions of ∼43 Å (assuming a Förster distance, *R*_0_, of ∼55 Å (Murphy et al., 2004)) (Figure S5A). In contrast, we observed a single non-zero peak centered at a significantly lower <E_FRET_> of ∼0.58 (*p*-value < 0.0005) for 30S IC-bound IF2(ppGpp), corresponding to a distance between our labeling positions of ∼52.5 Å, an increase of ∼9.5 Å relative to the ∼43 Å observed for IF2(GTP) (Figure S5B-D). Notably, the E_FRET_ distribution of 30S IC-bound IF2(ppGpp) closely resembles that of 30S IC-bound IF2(GDP), in that both distributions exhibit a single non-zero peak centered at an <E_FRET_> of ∼0.60. These results demonstrate that 30S IC-bound IF2(ppGpp) exhibits a conformation different from that of an IF2 that is active for rapid subunit joining (*i.e.*, IF2(GTP)) and similar to that of an IF2 that is inactive for rapid subunit joining (*i.e.*, IF2(GDP)).

### (p)ppGpp binding to IF2 mediates translational inhibition during transition phase

Our identification of an IF2 allele that is less sensitive to (p)ppGpp *in vitro* enabled us to test our initial hypothesis that accumulation of (p)ppGpp during growth reduces protein synthesis *in vivo* because (p)ppGpp binds IF2 and inhibits its function in translation. We generated a *B. subtilis* strain carrying a single copy of IF2 (*infB*) containing the double mutations (G226A, H230A) that *in vitro* reduce ppGpp binding without substantially inhibiting IF2 function (Figure 5C; Figure S4B). This strain grows equivalently to the wildtype parent throughout all phases of growth, validating that the mutant IF2 is functional *in vivo* (Figure S6A). We first tested how these mutations affect protein synthesis during late transition phase since in the wildtype background, protein synthesis is strongly inhibited in a subpopulation of cells during this period (Figure 1C). Mutations in IF2 that affect its binding to (p)ppGpp appear to significantly attenuate this phenotype (Figure 7 A, B; Figure S6 B, C). This attenuation is similar to that observed in the complete absence of (p)ppGpp (Figure 1C), suggesting it is due to the direct interaction of (p)ppGpp with IF2. Thus, these data indicate that during transition phase, (p)ppGpp binding to IF2 is sufficient to substantially inhibit translation.

**Figure 7.**
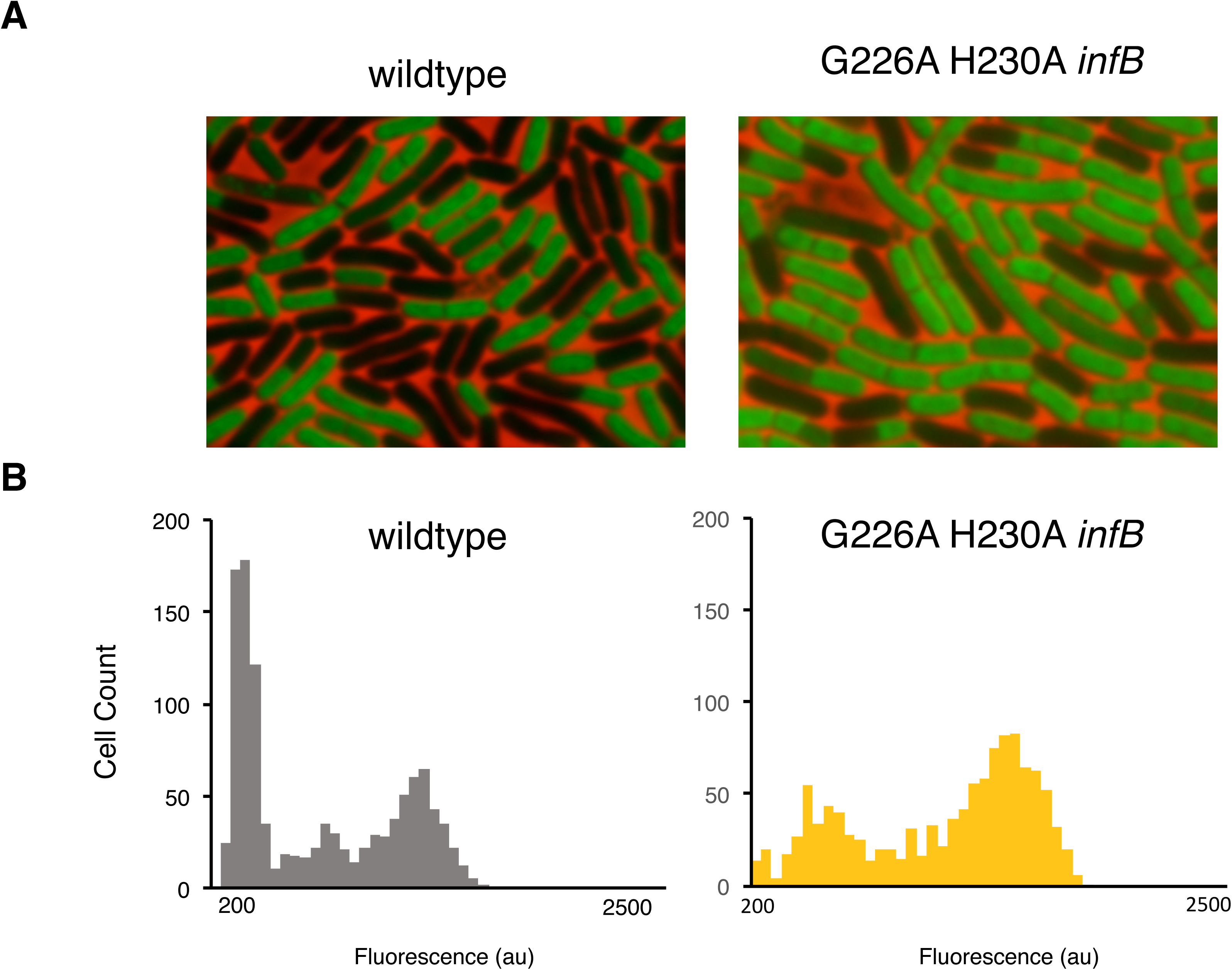
IF2 mediates inhibition of protein synthesis by (p)ppGpp during late transition phase. IF2 was validated as an *in vivo* target of (p)ppGpp by measuring protein synthesis in a strain expressing a G226A H230A double mutant IF2. **(A)** Representative pictures of WT (JDB 1772) and G226A H230A *infB* (JDB 4297) strains labelled with OPP during late transition phase. **(B)** Distributions of mean cell fluorescence of WT and G226A H230A *infB* strains during late transition phase. Late transition phase time point is the same as that in Figure 1. See also Figure S6

## Discussion

Here, we demonstrate that protein synthesis is actively attenuated in *B. subtilis* during the transition phase of growth, initially resulting in a subpopulation of cells that are deficient in protein synthesis. The alarmone (p)ppGpp is both necessary and sufficient for this phenomenon and acts through a mechanism that is, at least in part, a direct effect on the translational GTPase IF2. Thus, this regulatory mechanism of endogenous (p)ppGpp synthesis mediates the downregulation of the most energy-consuming process in cells as they enter quiescence.

Although the function of (p)ppGpp under basal (non-stringent) conditions has not been extensively investigated, loss of (p)ppGpp in *E. faecalis* leads to broad changes in physiology without stringent response activation (Gaca et al., 2013). More directly relevant to the present observations is the finding that the global translation rate per cell under basal conditions is elevated in a *S. elongatus* strain lacking the Rel (p)ppGpp synthetase (Puszynska and O’Shea, 2017). (p)ppGpp synthesis is also relevant to physiological situations including survival of pathogenic (Stapels et al., 2018) and commensal bacteria (Schofield et al., 2018). “Persisters” are rare bacterial cells in populations with increased tolerance to antibiotics that are thought to be characterized by relatively higher levels of (p)ppGpp (Hauryliuk et al., 2015). While the basis for this heterogeneity is not understood, an interesting question raised by our observations is whether the increased antibiotic tolerance of these cells is, at least in part, a consequence of the (p)ppGpp-dependent inhibition of protein synthesis.

We show that the (p)ppGpp sensitivity of IF2 can be altered by a specific double mutation in the G1 motif of the G domain, the site of (p)ppGpp binding (Milon et al., 2006). There are numerous examples of other ribosome associated GTPases that bind (p)ppGpp such as ObgE (Buglino et al., 2002; Feng et al., 2014; Persky et al., 2009), BipA (Kumar et al., 2015), RbgA (Corrigan et al., 2016; Pausch et al., 2018), HflX (Corrigan et al., 2016) and Era (Corrigan et al., 2016). Many of these reports demonstrate that (p)ppGpp binding affects their *in vitro* function in aspects of protein synthesis, such as ribosome assembly. A critical question, however, is assessing the implications of (p)ppGpp binding for the *in vivo* function of these proteins. Our demonstration that point mutations in IF2 affect (p)ppGpp binding and subsequent *in vitro* (Figure 5) and *in vivo* function (Figure 7) suggests that introducing similar mutations into the highly conserved G1 motif of other GTPases might be informative. Interestingly, a similar mutagenic strategy was reported very recently for the *E. coli* PurF glutamine amido-phoaphoribotransferease where a single point mutation yielded a protein that was insensitive to ppGpp inhibition (Wang et al., 2019).

Our studies also reveal the structural basis through which ppGpp targets IF2 to inhibit translation initiation. We find that ppGpp stabilizes a conformation of IF2 that is different from that which is stabilized by GTP, resulting in a reduced affinity of IF2(ppGpp) relative to IF2(GTP) for the 30S IC. Given that IF2 promotes the binding of fMet-tRNA^fMet^ to the 30S subunit during 30S IC assembly and subsequently accelerates subunit joining (Antoun et al., 2006), one way in which ppGpp may interfere with these activities is by decreasing the affinity of IF2 for the 30S IC and consequently reducing the fraction of 30S ICs harboring IF2. Notably, even when bound to the 30S IC, IF2(ppGpp) exhibits a conformation that differs from the conformation of IF2(GTP) that is active for rapid subunit joining. Thus, in addition to lowering the fraction of 30S ICs harboring IF2, our results indicate that ppGpp fails to allosterically activate 30S IC-bound IF2 for rapid subunit joining. Specifically, in the ppGpp-bound conformation of IF2, the distance between our labeling positions on dIV of IF2 and fMet-tRNA^fMet^ is ∼9.5 Å longer than in the GTP-bound conformation of IF2. These results suggest that dIV and the *N*-formyl-methionine and aminoacyl acceptor stem of fMet-tRNA^fMet^ are unable to form the same number and/or strength of stabilizing interactions in the presence of ppGpp that they make in the presence of GTP. Consequently, ppGpp may interfere with the formation and/or stabilization of the IF2-tRNA sub-complex on the 30S IC, the formation of which has been shown in structural studies to play a key role in stabilizing the binding and/or positioning of fMet-tRNA^fMet^ on the 30S IC (Julian et al., 2011; Simonetti et al., 2008). In our previous smFRET study, we demonstrated that recognition of the *N*-formyl-methionine and aminoacyl acceptor stem by dIV of IF2 was crucial for the allosteric activation of IF2(GTP), thereby enabling IF2 to catalyze rapid subunit joining (Caban et al., 2017). Thus, combined with the decreased affinity of IF2(ppGpp) for the 30S IC, our results suggest that the conformation of 30S IC-bound IF2(ppGpp) directly interferes with the ability of IF2 to promote rapid subunit joining to the 30S IC.

It is notable that the affinity of IF2(ppGpp) for the 30S IC and the conformation of 30S IC-bound IF2(ppGpp) are similar to those which we have reported for IF2(GDP) (Caban et al., 2017).Consistent with these similarities, Milón and colleagues have demonstrated that ppGpp and GDP establish the same hydrogen bonding network within the G-nucleotide binding pocket of IF2, resulting in very similar structures for the ppGpp- and GDP-bound G domains of IF2 (Milon et al., 2006). In terms of the activities of IF2, neither ppGpp or GDP are able to effectively stabilize the binding (Vinogradova et al., 2019) and/or positioning of fMet-tRNA^fMet^ on the 30S IC and catalyze rapid subunit joining. These similarities suggest that the mechanism through which ppGpp inhibits the activity of IF2 is through stabilizing the factor in an inactive conformation that is similar to that stabilized by GDP, thereby interfering with the activities of IF2 during translation initiation.

The present study raises several questions. First, what is the source of the observed heterogeneity of protein synthesis in single cells (Figure 1)? One possibility is heterogeneity in *sasA* and/or *sasB* expression resulting in different levels of (p)ppGpp across the population. Recently, it was observed that rare cells in exponential growth have high levels of *sasA* expression with concomitant physiological effects including induction of (p)ppGpp-dependent genes and enhanced antibiotic tolerance (Libby et al., 2019). However, the frequency of these cells in a population (∼1%) is much too low to account for the observed heterogeneity. The roughly bimodal character of the heterogeneity (Figure 1C) suggests that it is an example of bistability (Dubnau and Losick, 2006), a phenomenon often attributed to regulatory network architecture, specifically the presence of positive non-linear autoregulation or two mutually repressive repressors (Ferrell, 2002). It is therefore intriguing to note that SasB (but not SasA) is subject to positive allosteric regulation (Steinchen et al., 2018) which could generate a sharp threshold-like response such as has been characterized with *E. coli* RelA (Shyp et al., 2012). Furthermore, *sasB* is expressed during early transition phase (Tagami et al., 2012). Thus, the potential role of *sasB* in the heterogeneity of protein synthesis (Figure 1) will be the subject future investigation.

Second, is IF2 the only target of (p)ppGpp? The *in vitro* translation experiment using the double mutant IF2 (Figure 5D) suggests that there may be residual inhibition that is not IF2- dependent. One possible additional target is EF-Tu (Hamel and Cashel, 1974) and the *in vitro* affinity of EF-Tu for (p)ppGpp is similar to that of IF2 (Mitkevich et al., 2010). Consistent with EF-Tu being a target, residues in IF2 that are important for its (p)ppGpp sensitivity are conserved in EF-Tu. Thus, characterization of EF-Tu mutant alleles carrying mutations similar to the IF2 double mutant with reduced sensitivity to (p)ppGpp could provide insight of the role of EF-Tu in the down- regulation of protein synthesis during post-exponential growth.

Third, how is (p)ppGpp-mediated inhibition reversed? Conserved proteins capable of hydrolyzing (p)ppGpp have been identified in many bacterial species (Atkinson et al., 2011; Irving and Corrigan, 2018). In *B. subtilis*, the Rel protein is a bi-functional (p)ppGpp synthetase and hydrolase. Although the regulation of the synthetase activity is well understood, regulation of the hydrolase activity remains understudied. Binding of branched chain amino acids (BCAA) to the *Rhodobacter capsulatus* Rel protein stimulates (p)ppGpp hydrolysis (Fang and Bauer, 2018). Since the key residue important for BCAA binding is conserved in *B. subtilis* RelA, this mechanism could mediate (p)ppGpp hydrolysis during outgrowth. Furthermore, a nudix family pyrophosphatase from *Thermus thermophilus* involved in (p)ppGpp dependent growth control has been shown to degrade (p)ppGpp (Ooga et al., 2009). A *B. subtilis* homolog may therefore be necessary for the re-initiation of growth.

Although we have shown that (p)ppGpp-dependent inhibition of translational initiation attenuates protein synthesis during transition phase, what about other mechanisms of down- regulation? For example, some species exhibit a loss of ribosomes during extended periods of stationary phase (Deutscher, 2003; Piir et al., 2011) which would be expected to decrease overall protein synthesis. However, recent work suggests that ribosomes are actually in excess, and that only a fraction are active, especially under non-exponential growth (Dai et al., 2016; Li et al., 2018). This excess would be particularly useful for the exit from quiescence, since the ability to transition to maximal protein synthesis as quickly as possible would provide a clear selective advantage (Korem Kohanim et al., 2018; Remigi et al., 2019). From this perspective, a reversible process such as the competitive binding of an inhibitor such as (p)ppGpp to a translational GTPase would facilitate the necessary down-regulation of protein synthesis to minimize energy consumption without impairing an optimal response to growth-permissive conditions.

## Acknowledgements

We thank the members of our labs for helpful discussions and comments on the manuscript. SD was supported by NIH R01GM114213-03S1 and T32AI106711; JR and RG were supported by NIH R01GM084288 and R01GM119386, JD was supported by NIH R01GM114213, R21AI135427 and is a Burroughs-Welcome Investigator in the Pathogenesis of Infectious Diseases.

## Author Contributions

Conceived and designed the experiments: SD, JR, RG, and JD. Performed the experiments and analyzed the data: SD, KC and JR. Supervised the study: RG. and JD. Wrote the paper: SD, JR, KC, RG, and JD.

## STAR METHODS

### LEAD CONTACT AND MATERIALS AVAILABILITY

Further information and requests for resources and reagents should be directed to and will be fulfilled by the Lead Contact, Jonathan Dworkin

### EXPERIMENTAL MODEL AND SUBJECT DETAILS

#### Bacterial strains construction

Strain JDB 4295 was generated by sequential transformation of JDB 1772 with gDNA from JDB 4294 to delete ywaC then yjbM, followed by integration of pSD49. relA gene was deleted at the last step of construction via transformation with genomic DNA (gDNA) from JDB 4294. Strain JDB 4296 was generated by sequential transformation of JDB 1772 with gDNA from JDB 4294 to knockout ywaC then yjbM, followed by integration of pSD49 and then integration of pSD47. relA gene was deleted at the last step of construction via transformation with gDNA from JDB 4294. Strain JDB 4297 was generated using a standard transformation protocol with pSD54 at 37°C. Following transformation, cells were plated on selective media and grown overnight at 45°C. Two transformants were grown for approximately 8 hours in LB without selection at 25°C. These cultures were then diluted 1:10 and grown overnight in LB without selection at 25°C. Overnight cultures were then serially diluted and plated to isolate single colonies. Ten single colonies from each plate were then checked for sensitivity to the antibiotic originally used for selection. Two sensitive clones from each of the original cultures were isolated and correct integration of point mutations was confirmed by sequencing PCR reactions of the region of interest.

#### Plasmid construction

Plasmid pSD49 was constructed by amplifying B. subtilis ywaC from JDB 1772 gDNA using ywaC RBS SalI F and ywaC BamHI R and digested with SalI and BamHI then ligated to SalI BamHI digested pDR150. pSD47 was constructed by first amplifying the P_veg_ promoter from pVEG using P_veg_ BamHI F and P_veg_ HindIII R and digested with BamHI and HindIII. This was then ligated to BamHI HindIII digested pSac-cm to generate pSac-cm-P_veg_. The luc gene was amplified from pGL3 using luc RBS HindIII F and luc R EcoRI and digested using HindIII and EcoRI and ligated to pSac-cm-P_veg_. pSD54 was constructed by amplifying B. subtilis G226A infB using gDNA from JDB 1772. Mutations were added by overlap extension PCR using infB EcoRI F and infB G226A R to amplify the upper fragment and infB G226A F and infB BamHI R to amplify the lower fragment. B. subtilis G226A H230A infB was then amplified from B. subtilis G226A infB PCR. Mutations were added by overlap extension PCR using infB EcoRI F and infB H230A R to amplify the upper fragment and infB H230A F and infB BamHI R to amplify the lower fragment. This final PCR product was then digested using EcoRI and BamHI then ligated to EcoRI BamHI digested pMINIMAD2. pSD36 was constructed by amplifying B. subtilis infB from JDB 1772 gDNA using infB NdeI F and infB BamHI R and digested with NdeI and BamHI and ligated to NdeI BamHI digested pETPHOS. pSD53 was constructed by amplifying B. subtilis G226A H230A infB from pSD54 using infB NdeI F and infB BamHI and digested with NdeI and BamHI and ligated to NdeI BamHI digested pETPHOS. pSD30 was constructed by amplifying B. subtilis fusA from JDB 1772 gDNA using fusA NdeI F and fusA BamHI R and digested with NdeI and BamHI and ligated to NdeI BamHI digested pETPHOS. pSD56 was constructed by amplifying E. coli 1-455 relA from JDE 1497 gDNA using E. coli relA NdeI F and E. coli relA BamHI R and digested with NdeI and BamHI and ligated to NdeI BamHI digested pETPHOS.

### METHOD DETAILS

#### Growth curves

Growth curves were performed in a Tecan Infinite m200 plate reader at 37 °C with continuous shaking and OD_600_ measurements were made every five minutes. Cultures were grown from single colonies from fresh LB plates grown overnight at 37 °C. Exponential phase starter cultures (OD_600_ ∼ 0.5-1.0) were diluted to OD_600_ = 0.01 and grown in 96-well Nunclon Delta surface clear plates (Thermo Scientific) with 150 μL per well. All growth curves were done in triplicate and media-only wells were used to subtract background absorbance.

#### OPP labeling

Click-iT Plus OPP Protein Synthesis Assay Kit (Invitrogen) was used to label cells with OPP following manufacturer’s instructions. 450 μL of cells at given time points were transferred to disposable glass tubes. OPP was added to a final concentration of 13 μM. Labelling was performed at 37 °C on a roller drum for 20 min and all subsequent steps were done at RT. Cells were harvested by centrifugation at 16K RCF for 1 min and re-suspended in 100 μL of 3.7% formaldehyde in PBS for fixation. Cells were fixed for 10 min, harvested, and permeabilized using 100 μL of 0.5% Triton X-100 in PBS for 15 min. Cells were labelled using 100 μL of 1X Click-iT cocktail for 20 min in the dark. Cells were harvested and washed one time using Click-iT rinse buffer and then re-suspended in 20-40 μL of PBS for imaging or in 150 μL of PBS for fluorescence measurement on a Tecan Infinite m200 plate reader in 96-well flat bottom White sided plates (Greiner Bio-One). Images were analyzed using Image J.

#### Luminescence growth curves

Cultures were grown in LB from single colonies grown overnight at 37 °C on LB plates. Cultures in exponential phase (OD_600_ ∼ 0.5-1.0) were diluted to OD_600_ = 0.1 in 150 μL LB containing 4.7 mM D-luciferin (Goldbio) and grown in a 96-well flat bottom white sided plates (Greiner Bio-One) plates in triplicate. OD_600_ and luminescence measurements were made every five min using a Tecan Infinite m200 plate reader and media only wells were used to subtract background.

#### RNA quantification

RNA was quantified from cultures grown in LB as above. At given time points 14 mL of the cultures were pelleted at 8 K RCF for 10 minutes at room temperature and frozen at −80 °C. Pellets were re-suspended in TRIzol (Invitrogen) to match based on OD_600_. ∼ 5 OD_600_ units of all cultures were lysed using a FastPrep 24 5G (MP Biomedicals). Lysates were spun down at 20 K RCF for 20 min and RNA was extracted using the Direct-Zol RNA miniprep kit (Zymo Research). RNA samples were DNAse I treated following manufactures protocol (NEB) and 1 μg of RNA was used to generate cDNA using the High Capacity cDNA Reverse Transcription Kit (Applied Biosystems). cDNAs were diluted 1:200 and used as templates for qPCR. qPCRs were preformed using SYBR green. Primers were design using the PrimeQuest Tool (IDT). No cDNA and no RT controls were used to ensure signal was specific to desired RNAs.

#### *in vitro* translation assays

Translation assays used the PURExpress system (NEB) following the manufacture’s protocol and a plasmid encoding a CotE-FLAG fusion protein as template DNA (Pereira et al., 2015). ppGpp (TriLink Biotechnologies) at the specified concentrations was added to translation reactions. Reactions were run for 20 min each at 37 °C and stopped by adding 2X SDS loading buffer. Synthesized proteins were separated by SDS-PAGE, transferred to PVDF membranes and visualized by probing with an anti-FLAG HRP antibody (Sigma). Mutant IF2 was assayed using a ΔIF123 PURExpress kit (NEB) supplemented with equal concentrations of purified *E. coli* IF1 and IF3. Reactions were run essentially as above but 0.47 μM of either WT or mutant *B. subtilis* IF2 was added to each reaction as the sole source of IF2. WT and mutant IF2s were purified as previously described (Fei et al., 2010). Band intensities were analyzed using ImageJ.

#### DRaCALA binding assays

Radiolabeled (p)ppGpp was generated essentially as described (Corrigan et al., 2016). Briefly, purified *E. coli* RelA N-terminal mutant protein (amino acids 1-455) was incubated overnight at 30°C with [α-^32^P]- GTP (PerkinElmer) in 50 mM Tris (pH 8), 15 mM MgOAc, 60 mM KOAc, 30 mM NH4OAc, 0.2 mM EDTA, and 0.5 mM PMSF. Reactions were supplemented with 8 mM cold ATP. Conversion of GTP to (p)ppGpp (>90%) was monitored by thin layer chromatography on PEI- cellulose plates in 1.5 M KH2PO4 (pH 3.6). DRaCALA binding assays were carried out essentially as described (Corrigan et al., 2016; Roelofs et al., 2011). 6 μM protein was incubated with 55.5 nM [α-^32^P]-labeled (p)ppGpp in 40 mM Tris (pH 8), 100 mM NaCl, 10 mM MgCl_2_, and 2mM PMSF. Reactions were incubated for 5 min at RT and 2.5 μL of each reaction was spotted onto nitrocellulose membranes and dried completely at RT. Spots were exposed for 30 min on a phosphor storage screen and visualized (GE Typhoon). Inner and outer ring intensities were quantified using ImageJ. Reactions where protein was not added were used to subtract background.

#### smFRET experiments

All of the *E. coli* components for assembling 30S ICs, including 30S ribosomal subunits, 5’- biotinylated mRNA, Cyanine (Cy) 3-labeled fMet-tRNA^fMet^ (labeled with maleimide-derivatized Cy3 at the naturally occurring 4-thiouridine at residue position 8), IF1, and Cy5-labeled IF2 (labeled with maleimide-derivatized Cy5 at an engineered cysteine at residue position 810) were prepared as previously described (Caban et al., 2017). 30S ICs lacking IF2 and IF3 were assembled by combining 0.6 μM 30S subunits, 1.8 μM 5’-biotinylated mRNA, 0.8 μM Cy3-labeled fMet-tRNA^fMet^, and 0.9 μM IF1 in Tris-Polymix Buffer (50 mM Tris-OAc (pH_RT_ = 7.5), 100 mM KCl, 5 mM NH_4_OAc, 5 mM Mg(OAc)_2_, 0.1 mM EDTA, 5 mM putrescine-HCl, 1 mM spermidine-free base, and 6 mM β- mercaptoethanol). The reaction was incubated at 37 °C for 10 minutes then on ice for an additional 5 minutes. Small aliquots of 30S ICs were flash-frozen in liquid nitrogen and stored at –80 °C.

To conduct smFRET experiments, 30S ICs assembled were first diluted to a final concentration of 75 pM in the presence of 2 uM IF1, 25 nM Cy5-labeled IF2, and 1 mM GTP or ppGpp. 30S ICs were then tethered to the polyethylene glycol (PEG)/biotin-PEG-derivatized surface of a microfluidic observation flowcell using a biotin-streptavidin-biotin between the 5’- biotinylated mRNA and the biotin-PEG. Untethered 30S ICs were flushed from the flowcell, and tethered 30S ICs were buffer exchanged, by washing the flowcell with Imaging Buffer (Tris-Polymix Buffer with an oxygen scavenging system composed of 2.5 mM 3,4-dihydroxybenzoic acid (PCA) and 250 mM protocatechuate 3,4-dioxygenase (PCD) and a triplet-state quencher cocktail composed of 8.4 mM 1,3,5,7-cyclooctatetraene (COT) and 8.7 mM 3-nitrobenzyl alcohol (NBA)) supplemented with 2 uM IF1, 25 nM Cy5-labeled IF2, and 1 mM GTP or ppGpp in order to enable rebinding of these components to 30SICs from which they might dissociates during the course of imaging. Finally, 30S ICs were imaged at single-molecule resolution and at a 0.1 sec per frame acquisition time using a laboratory-built, prism-based total internal reflection fluorescence (TIRF) microscope as previously described (Caban et al., 2017). A previously described approach (Desai and Gonzalez, 2019) was used to identify fluorophores and classify them into ‘fluorophore’ or ‘background’ classes; align the Cy3 and Cy5 imaging channels; fit individual Cy3 and Cy5 fluorophore to 2D Gaussians and estimate and, in the case of Cy5, bleedthrough correct the Cy3 and Cy5 fluorescence emission intensity *versus* time trajectories; and generate the E_FRET_ *versus* time trajectories. Only those trajectories exhibiting a signal-to-background (SBR) of 3.5:1 or greater as well as single-step photobleaching of Cy3 within the observation time were selected for further analyses.

In order to estimate the rate constants for the association of IF2 with the 30S IC (*k*_a_) and for the dissociation of IF2 from the 30S IC (*k*_d_) we began by estimating a ‘consensus’ hidden Markov model (HMM) of the E_FRET_ *versus* time trajectories using a slight extension of the variational Bayes approach we introduced in the vbFRET algorithm (Bronson et al., 2009) Briefly, instead of using a likelihood function for each E_FRET_ *versus* time trajectory, we used a single likelihood function that simultaneously includes all of the E_FRET_ *versus* time trajectories in a dataset to arrive at a log-likelihood function given by

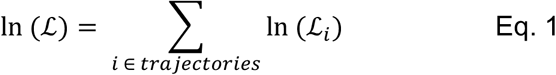

where ℒ*_i_* is the variational approximation of the likelihood function for a single trajectory. A further development of this approach in a hierarchical context underlies the hFRET algorithm that we have recently reported(Hon and Gonzalez, 2019). Using this consensus HMM approach, we estimated HMMs for 1-6 states and performed model selection using the highest evidence lower bound (ELBO) as described in Bronson et al., 2009. In all cases, the 2-state HMM yielded the highest ELBO. The transition matrix obtained from this 2-state model consists of a 2 x 2 matrix in which the off-diagonal elements correspond to the number of times a transition takes place between the IF2-free and the IF2-bound states of the 30S IC and the on-diagonal elements correspond to the number of times a transition does not take place. The 2 rows of this matrix parameterize Dirichlet distributions and, for each Dirichlet distribution, we calculated the lower bounds (2.5 %) and upper bounds (97.5 %) of the transition probability using the inverse cumulative distribution function of the corresponding Dirichlet distribution. These transition probabilities (*p*) were used to calculate rate constants (*k*) using the equation

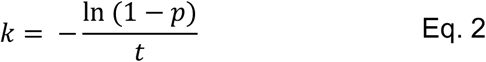

where *t* is the time between successive data points (*i.e.*, the acquisition time) (*t* = 0.1 sec). Finally, we calculated *k*_a_ using the equation

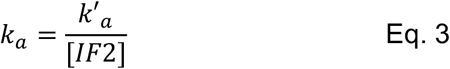

where *k*’_a_ is the pseudo-first-order association rate constant calculated using Eq. 2 and [IF2] is the concentration of IF2, and we calculated *k*_d_ directly from Eq 2. The equilibrium dissociation constant for IF2 binding to the 30S IS (*K*_d_) was obtained by summing the columns of the 2 x 2 transition matrix to obtain the total number of data points in which the 30S IC was either in the IF2-free state or the IF2-bound state. These sums can then be used to parameterize a Dirichlet distribution describing the fraction of 30S ICs in the IF2-bound state (*f*_b_). The lower bounds and upper bounds of *f*_b_ were calculated using the inverse cumulative distribution function of this Dirichlet distribution, as described above, and the *K*_d_ was calculated using the equation *K*_d_ / [IF2] = (1/*f*_b_) – 1.

### QUANTIFICATION AND STATISTICAL ANALYSIS

Western blots, DRaCALA images, and cell fluorescence intensities were quantified using ImageJ. Statistical significance was determined using an unpaired two-tailed Student’s t test unless otherwise stated.

---

**Figure S1.**
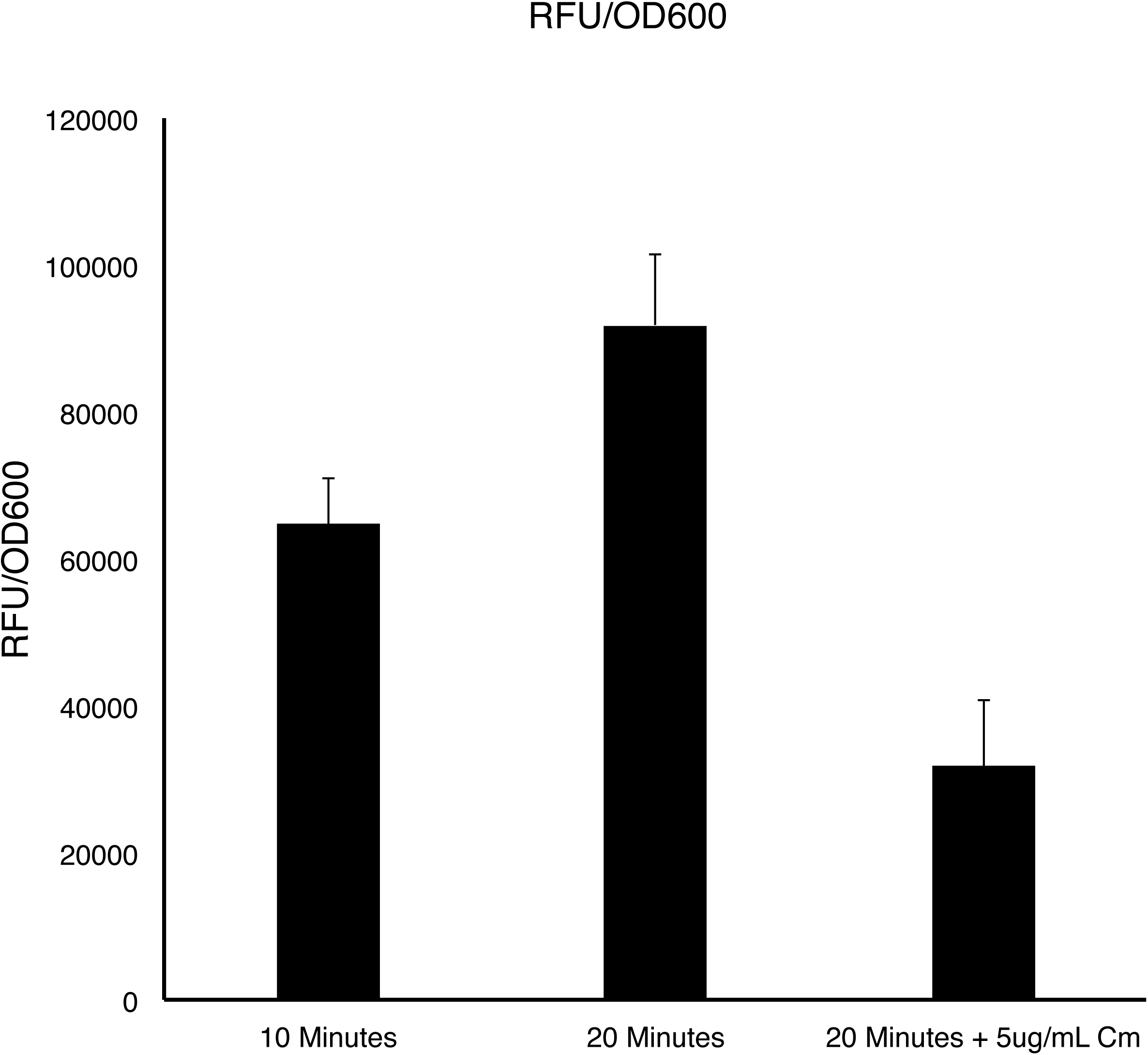
OPP does not arrest protein synthesis (Related to Figure 1) The effect of OPP addition on protein synthesis was tested during exponential phase. OPP was added to exponentially growing cultures of WT *B. subtilis* and total fluorescence was measured at 10 minutes and 20 minutes post OPP addition. Increased fluorescence was detected at the 20 minute time point compared to 10 minute time point indicating continued protein synthesis in the presence of OPP. Chloramphenicol was added to a separate culture in combination with OPP. Decreased fluorescence indicates that OPP is sensitive to translational inhibition (means ± SDs).

**Figure S2.**
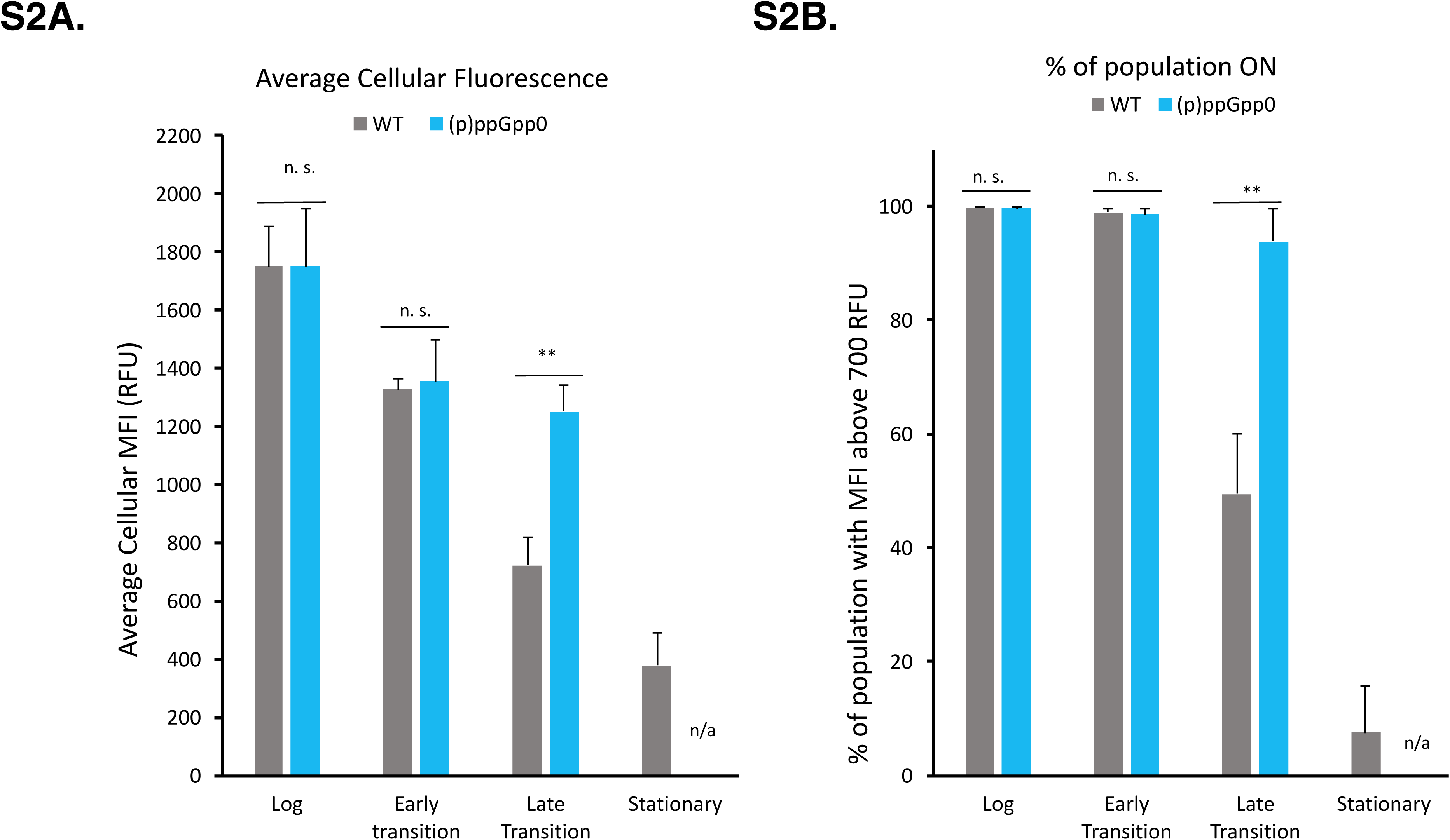
Average cellular fluorescence and % of population “ON” throughout growth in WT and (p)ppGpp null strains (Related to Figure 1) (A) Average cell fluorescence and (B) % of population ON were quantified from ∼1400 cells in three separate experiments (means ± SDs). n.s. p > 0.05, *p < 0.05, **p < 0.01, ***p < 0.001

**Figure S3.**
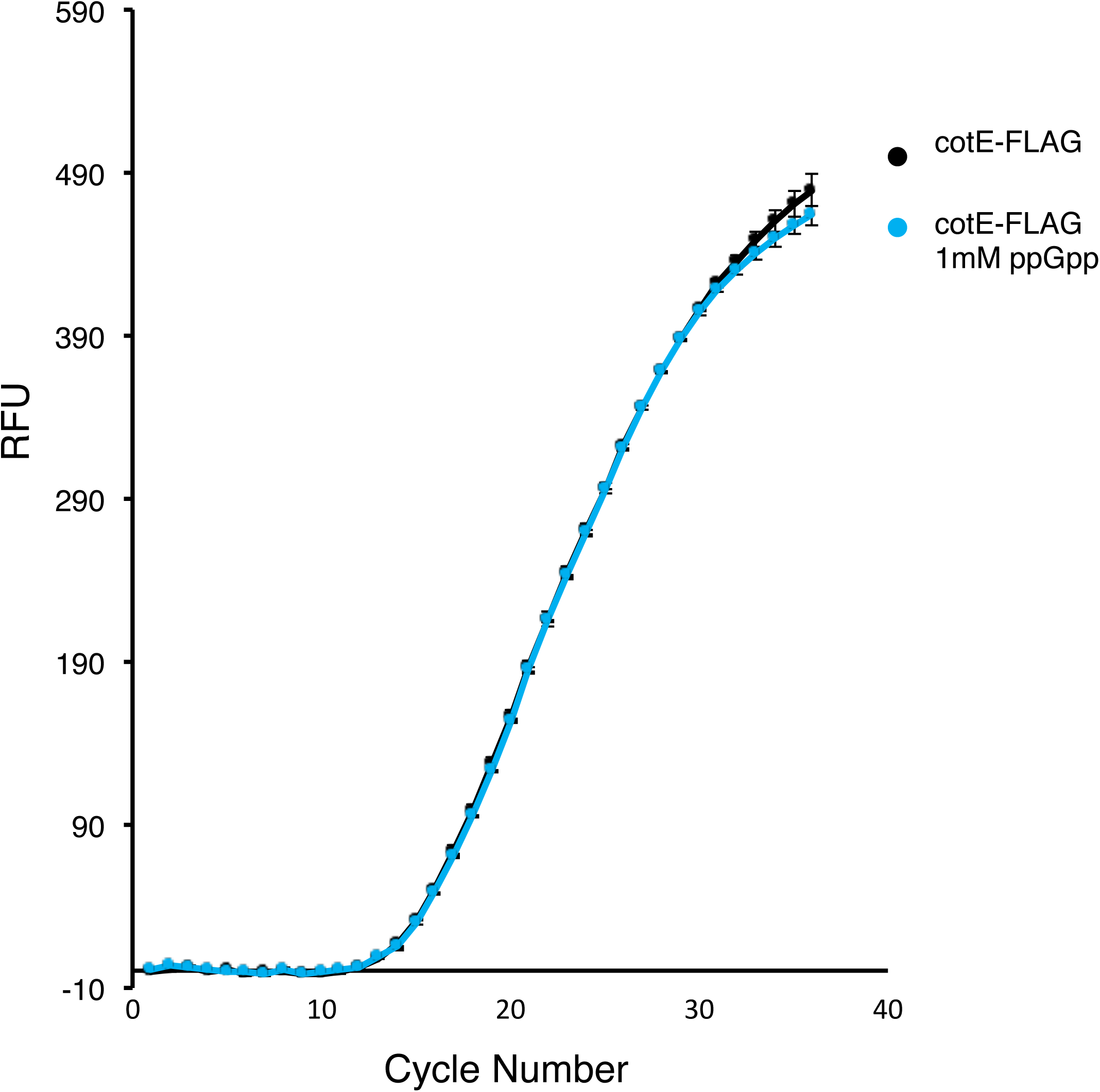
RT-qPCR of *in vitro* transcribed mRNA in the presence of ppGpp (Related to Figure 4) RNA synthesis was measured by quantifying total mRNA produced in the presence of a concentration of ppGpp (1mM) that significantly inhibits protein production. RNA was quantified using RT-qPCR (means ± SDs). Compare with 1mM concentration in Figure 4.

**Figure S4.**
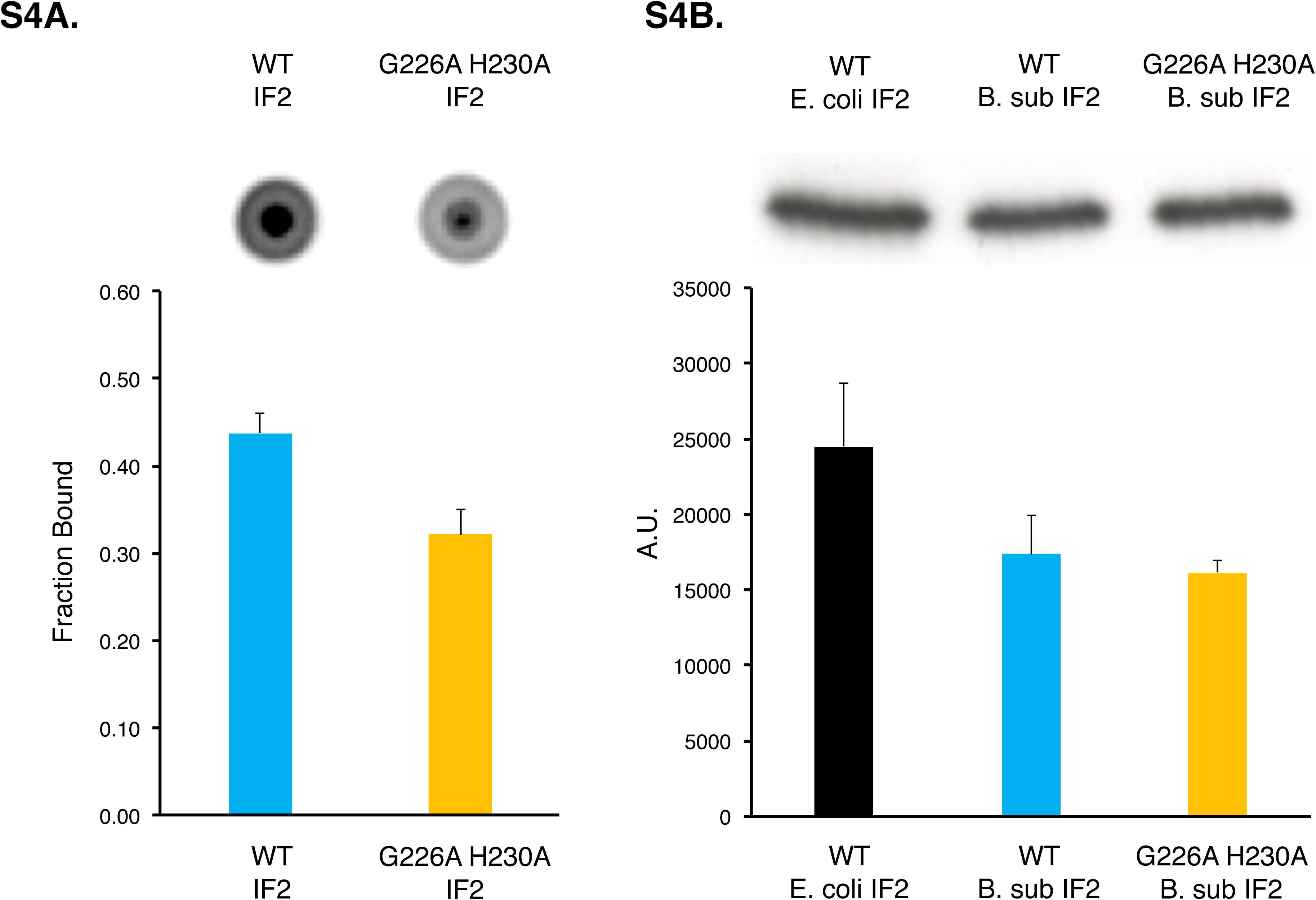
G226A H230A IF2 mutant is inhibited in binding GTP but not in function (Related to Figure 5) (A) GTP binding and (B) function of G226A H230A IF2 mutant was assayed using DRaCALA assay and an *in vitro* transcription-translation assay respectively (means ± SDs).

**Figure S5.**
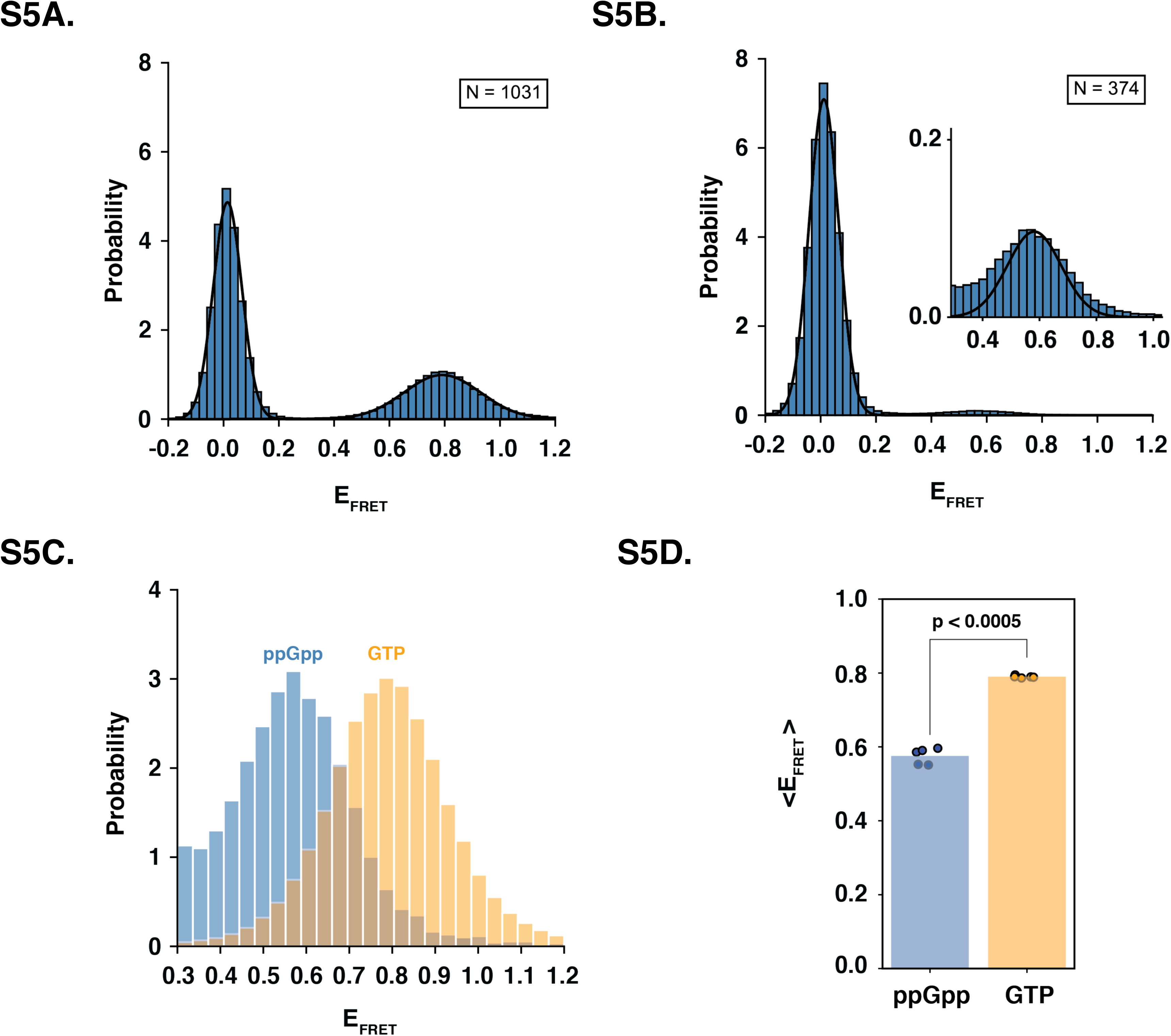
IF2(GTP) and IF2(ppGpp) exhibit distinct conformations when bound to the 30S IC (Related to Figure 6) One-dimensional histograms of E_FRET_ value distributions corresponding to the interaction of IF2 with the 30S IC for **(A)** IF2(GTP) and **(B**) IF2(ppGpp). Both histograms were fitted to a two-Gaussian mixture model in which the Gaussian centered at the zero E_FRET_ value corresponds to the IF2-free state of the 30S IC and the Gaussian centered at the non-zero E_FRET_ value corresponds to the IF2-bound state of the 30S IC. The center of each fitted Gaussian was used to determine the mean E_FRET_ value for the corresponding state (<E_FRET_>). The Gaussians corresponding to the IF2-free, IF2(GTP)-bound, and IF2(ppGpp)-bound states of the 30S IC had <E_FRET_>s of ∼0.0 (in both histograms), ∼0.80, and ∼0.58, respectively. **(C)** To qualitatively compare the E_FRET_ distributions for the IF2(GTP)-bound (orange) and IF2(ppGpp)-bound (blue) states of the 30S IC, the portion of the distributions with E_FRET_ values greater than 0.3 were normalized such that the areas of the two distributions were equivalent. Although there is some overlap between the distributions, the two distributions exhibit distinct <E_FRET_>s. **(D)** To quantitatively demonstrate that the difference between the <E_FRET_>s of the two samples (*i.e.*, the IF2(GTP)-bound state of the 30S IC and the IF2(ppGpp)-bound states of the 30S IC) is statistically meaningful, we randomly separated all of the E_FRET_ *versus* time trajectories for each sample into five groups, calculated the <E_FRET_> for each group in each sample, and conducted a two-sample t-test, obtaining a p-value < 0.0005.

**Figure S6.**
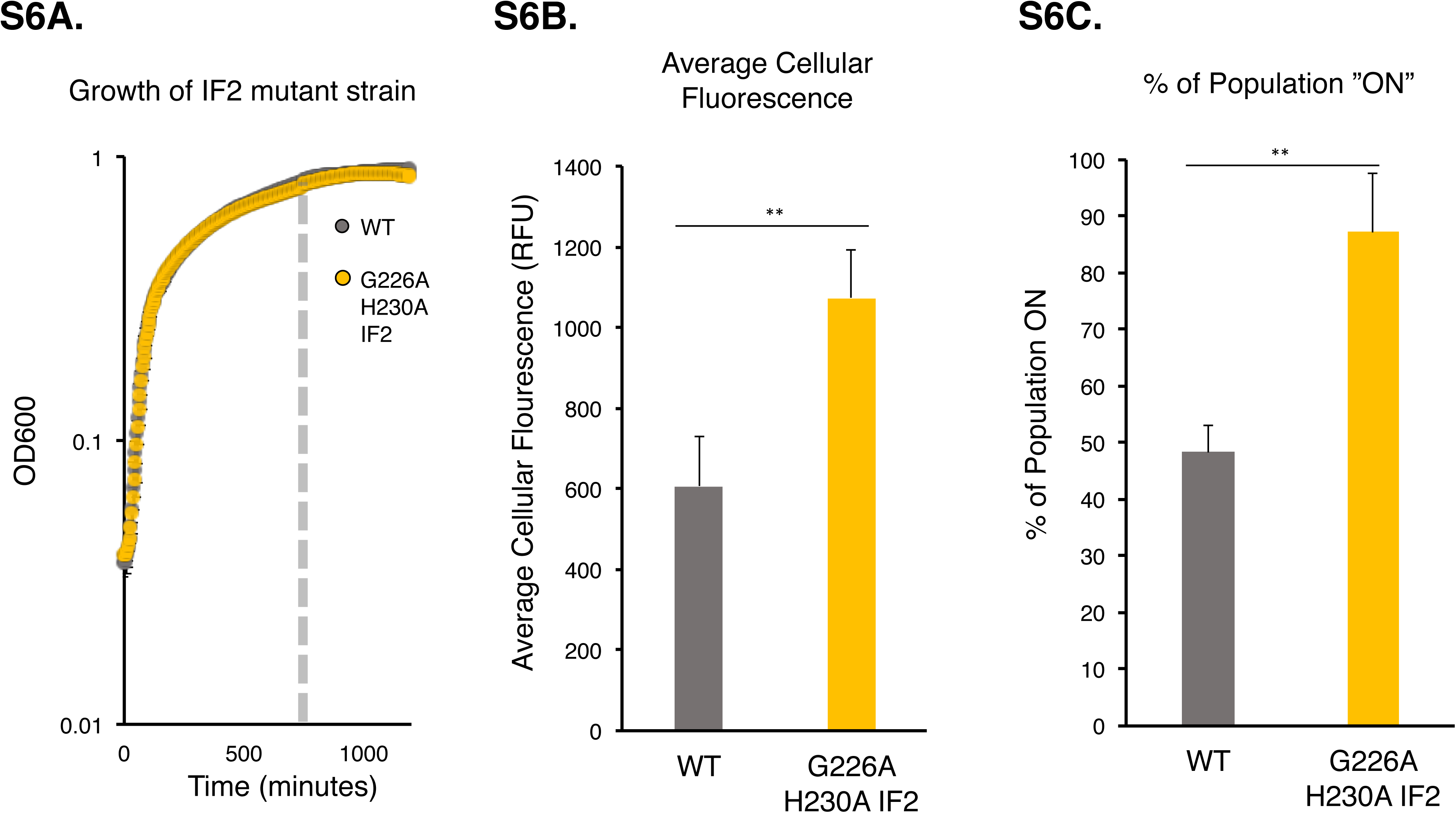
Growth curve average cellular fluorescence and % of population ON throughout growth in WT and G226A H230A *infB* strains (Related to Figure 7) Growth curve of G226A H230A IF2 strain is equivalent to WT. (B) Average cell fluorescence and (C) % of population ON were quantified from ∼1400 cells in three separate experiments (means ± SDs). n.s. p > 0.05, *p < 0.05, **p < 0.01, ***p < 0.001

**Table S1.** Association rate constant (k_a_), dissociation rate constant (k_d_), and equilibrium dissociation constant (K_d_) for the interaction of IF2(GTP) and IF2(ppGpp) with the 30S IC (Related to Figure 6) ^a^Values correspond to 95% credible interval ranges calculated as described in Methods.

| | $k_a$ ($\mu\text{M}^{-1} \text{s}^{-1}$)<sup>a</sup> | $k_d$ ($\text{s}^{-1}$)<sup>a</sup> | $K_d$ (nM)<sup>a</sup> |
| --- | --- | --- | --- |
| IF2(GTP) | 1.56 ± 0.08 | 0.091 ± 0.004 | 45 ± 0.2 |
| IF2(ppGpp) | 2.5 ± 0.1 | 1.45 ± 0.07 | 534 ± 10 |

## References

Antoun, A., Pavlov, M.Y., Andersson, K., Tenson, T., and Ehrenberg, M. (2003). The roles of initiation factor 2 and guanosine triphosphate in initiation of protein synthesis. EMBO J 22, 5593–5601.

Antoun, A., Pavlov, M.Y., Lovmar, M., and Ehrenberg, M. (2006). How initiation factors tune the rate of initiation of protein synthesis in bacteria. EMBO J 25, 2539–2550.

Antoun, A., Pavlov, M.Y., Tenson, T., and Ehrenberg, M.M. (2004). Ribosome formation from subunits studied by stopped-flow and Rayleigh light scattering. Biol Proced Online 6, 35–54.

Arenz, S., Abdelshahid, M., Sohmen, D., Payoe, R., Starosta, A.L., Berninghausen, O., Hauryliuk, V., Beckmann, R., and Wilson, D.N. (2016). The stringent factor RelA adopts an open conformation on the ribosome to stimulate ppGpp synthesis. Nucleic Acids Res 44, 6471–6481.

Artsimovitch, I., Patlan, V., Sekine, S., Vassylyeva, M.N., Hosaka, T., Ochi, K., Yokoyama, S., and Vassylyev, D.G. (2004). Structural basis for transcription regulation by alarmone ppGpp. Cell 117, 299–310.

Atkinson, G.C., Tenson, T., and Hauryliuk, V. (2011). The RelA/SpoT homolog (RSH) superfamily: distribution and functional evolution of ppGpp synthetases and hydrolases across the tree of life. PLoS One 6, e23479.

Boes, N., Schreiber, K., and Schobert, M. (2008). SpoT-triggered stringent response controls usp gene expression in Pseudomonas aeruginosa. J Bacteriol 190, 7189–7199.

Bronson, J.E., Fei, J., Hofman, J.M., Gonzalez, R.L., Jr., and Wiggins, C.H. (2009). Learning rates and states from biophysical time series: a Bayesian approach to model selection and single-molecule FRET data. Biophys J 97, 3196–3205.

Brown, A., Fernandez, I.S., Gordiyenko, Y., and Ramakrishnan, V. (2016). Ribosome-dependent activation of stringent control. Nature 534, 277–280.

Buckstein, M.H., He, J., and Rubin, H. (2008). Characterization of nucleotide pools as a function of physiological state in Escherichia coli. J Bacteriol 190, 718–726.

Buglino, J., Shen, V., Hakimian, P., and Lima, C.D. (2002). Structural and biochemical analysis of the Obg GTP binding protein. Structure 10, 1581–1592.

Caban, K., Pavlov, M., Ehrenberg, M., and Gonzalez, R.L., Jr. (2017). A conformational switch in initiation factor 2 controls the fidelity of translation initiation in bacteria. Nat Commun 8, 1475.

Corrigan, R.M., Bellows, L.E., Wood, A., and Grundling, A. (2016). ppGpp negatively impacts ribosome assembly affecting growth and antimicrobial tolerance in Gram-positive bacteria. Proc Natl Acad Sci U S A 113, E1710–1719.

Dai, X., Zhu, M., Warren, M., Balakrishnan, R., Patsalo, V., Okano, H., Williamson, J.R., Fredrick, K., Wang, Y.P., and Hwa, T. (2016). Reduction of translating ribosomes enables Escherichia coli to maintain elongation rates during slow growth. Nat Microbiol 2, 16231.

Desai, B.J., and Gonzalez, R.L. (2019).

Deutscher, M.P. (2003). Degradation of stable RNA in bacteria. J Biol Chem 278, 45041–45044.

Dubnau, D., and Losick, R. (2006). Bistability in bacteria. Molecular microbiology 61, 564–572.

Fang, M., and Bauer, C.E. (2018). Regulation of stringent factor by branched-chain amino acids. Proc Natl Acad Sci U S A 115, 6446–6451.

Fei, J., Wang, J., Sternberg, S.H., MacDougall, D.D., Elvekrog, M.M., Pulukkunat, D.K., Englander, M.T., and Gonzalez, R.L. (2010). A Highly Purified, Fluorescently Labeled In Vitro Translation System for Single-Molecule Studies of Protein Synthesis. In Single Molecule Tools: Fluorescence Based Approaches, Part A, pp. 221–259.

Feng, B., Mandava, C.S., Guo, Q., Wang, J., Cao, W., Li, N., Zhang, Y., Zhang, Y., Wang, Z., Wu, J., et al. (2014). Structural and functional insights into the mode of action of a universally conserved Obg GTPase. PLoS Biol 12, e1001866.

Ferrell, J.E., Jr. (2002). Self-perpetuating states in signal transduction: positive feedback, double-negative feedback and bistability. Curr Opin Cell Biol 14, 140–148.

Friesen, J.D., and Fiil, N. (1973). Accumulation of Guanosine Tetraphosphate in T7 Bacteriophage-Infected Escherichia coli J Bacteriol 113, 697–703.

Gaca, A.O., Colomer-Winter, C., and Lemos, J.A. (2015). Many means to a common end: the intricacies of (p)ppGpp metabolism and its control of bacterial homeostasis. J Bacteriol 197, 1146–1156.

Gaca, A.O., Kajfasz, J.K., Miller, J.H., Liu, K., Wang, J.D., Abranches, J., and Lemos, J.A. (2013). Basal levels of (p)ppGpp in Enterococcus faecalis: the magic beyond the stringent response. MBio 4, e00646–00613.

Hamel, E., and Cashel, M. (1974). Guanine nucleotides in protein synthesis. Utilization of pppGpp and dGTP by initiation factor 2 and elongation factor Tu. Archives of biochemistry and biophysics 162, 293–300.

Haseltine, W.A., and Block, R. (1973). Synthesis of guanosine tetra- and pentaphosphate requires the presence of a codon-specific, uncharged transfer ribonucleic acid in the acceptor site of ribosomes. Proc Natl Acad Sci U S A 70, 1564–1568.

Hauryliuk, V., Atkinson, G.C., Murakami, K.S., Tenson, T., and Gerdes, K. (2015). Recent functional insights into the role of (p)ppGpp in bacterial physiology. Nat Rev Microbiol 13, 298–309.

Hinnebusch, A.G. (2005). Translational regulation of GCN4 and the general amino acid control of yeast. Annu Rev Microbiol 59, 407–450.

Hon, J., and Gonzalez, R.L., Jr. (2019). Bayesian-Estimated Hierarchical HMMs Enable Robust Analysis of Single-Molecule Kinetic Heterogeneity. Biophys J 116, 1790–1802.

Irving, S.E., and Corrigan, R.M. (2018). Triggering the stringent response: signals responsible for activating (p)ppGpp synthesis in bacteria. Microbiology 164, 268–276.

Julian, P., Milon, P., Agirrezabala, X., Lasso, G., Gil, D., Rodnina, M.V., and Valle, M. (2011). The Cryo-EM structure of a complete 30S translation initiation complex from Escherichia coli. PLoS Biol 9, e1001095.

Korem Kohanim, Y., Levi, D., Jona, G., Towbin, B.D., Bren, A., and Alon, U. (2018). A Bacterial Growth Law out of Steady State. Cell Rep 23, 2891–2900.

Krasny, L., and Gourse, R.L. (2004). An alternative strategy for bacterial ribosome synthesis- Bacillus subtilis rRNA transcription regulation. EMBO 23, 4473–4483.

Kriel, A., Bittner, A.N., Kim, S.H., Liu, K., Tehranchi, A.K., Zou, W.Y., Rendon, S., Chen, R., Tu, B.P., and Wang, J.D. (2012). Direct regulation of GTP homeostasis by (p)ppGpp: a critical component of viability and stress resistance. Mol Cell 48, 231–241.

Kriel, A., Brinsmade, S.R., Tse, J.L., Tehranchi, A.K., Bittner, A.N., Sonenshein, A.L., and Wang, J.D. (2014). GTP dysregulation in Bacillus subtilis cells lacking (p)ppGpp results in phenotypic amino acid auxotrophy and failure to adapt to nutrient downshift and regulate biosynthesis genes. J Bacteriol 196, 189–201.

Kumar, V., Chen, Y., Ero, R., Ahmed, T., Tan, J., Li, Z., Wong, A.S., Bhushan, S., and Gao, Y.G. (2015). Structure of BipA in GTP form bound to the ratcheted ribosome. Proc Natl Acad Sci U S A 112, 10944–10949.

Legault, L., Jeantet, C., and Gros, F. (1972). Inhibition of in vitro protein synthesis by ppGpp. FEBS Lett 27, 71–75.

Lennon, J.T., and Jones, S.E. (2011). Microbial seed banks: the ecological and evolutionary implications of dormancy. Nat Rev Microbiol 9, 119–130.

Li, S.H., Li, Z., Park, J.O., King, C.G., Rabinowitz, J.D., Wingreen, N.S., and Gitai, Z. (2018). Escherichia coli translation strategies differ across carbon, nitrogen and phosphorus limitation conditions. Nat Microbiol 3, 939–947.

Libby, E.A., Reuveni, S., and Dworkin, J. (2019). Multisite phosphorylation regulates phenotypic variability in antibiotic tolerance. Nature Communications accepted.

Lindahl, L., Post, L., and Nomura, M. (1976). DNA-dependent in vitro synthesis of fibosomal proteins, protein elongation factors, and RNA polymerase subunit alpha: inhibition by ppGpp. Cell 9, 439–448.

Liu, J., Xu, Y., Stoleru, D., and Salic, A. (2012). Imaging protein synthesis in cells and tissues with an alkyne analog of puromycin. Proc Natl Acad Sci U S A 109, 413–418.

Liu, K., Bittner, A.N., and Wang, J.D. (2015). Diversity in (p)ppGpp metabolism and effectors. Curr Opin Microbiol 24, 72–79.

Loveland, A.B., Bah, E., Madireddy, R., Zhang, Y., Brilot, A.F., Grigorieff, N., and Korostelev, A.A. (2016). Ribosome*RelA structures reveal the mechanism of stringent response activation. Elife 5.

Maracci, C., and Rodnina, M.V. (2016). Review: Translational GTPases. Biopolymers 105, 463–475.

Milon, P., Tischenko, E., Tomsic, J., Caserta, E., Folkers, G., Teana, A.L., Rodnina, M.V., Pon, C.L., Boelens, R., and Gualerzi, C.O. (2006). The nucleotide-binding site of bacterial translation initiation factor 2 (IF2) as a metabolic sensor. Proc Natl Acad Sci U S A 103, 13962–13967.

Mirouze, N., Prepiak, P., and Dubnau, D. (2011). Fluctuations in spo0A transcription control rare developmental transitions in Bacillus subtilis. PLoS Genet 7, e1002048.

Mitkevich, V.A., Ermakov, A., Kulikova, A.A., Tankov, S., Shyp, V., Soosaar, A., Tenson, T., Makarov, A.A., Ehrenberg, M., and Hauryliuk, V. (2010). Thermodynamic characterization of ppGpp binding to EF-G or IF2 and of initiator tRNA binding to free IF2 in the presence of GDP, GTP, or ppGpp. Journal of molecular biology 402, 838–846.

Murphy, M.C., Rasnik, I., Cheng, W., Lohman, T.M., and Ha, T. (2004). Probing single-stranded DNA conformational flexibility using fluorescence spectroscopy Biophysical Journal 86, 2530–2537.

Murray, H.D., Schneider, D.A., and Gourse, R.L. (2003). Control of rRNA expression by small molecules is dynamic and nonredundant. Mol Cell 12, 125–134.

Nanamiya, H., Kasai, K., Nozawa, A., Yun, C.S., Narisawa, T., Murakami, K., Natori, Y., Kawamura, F., and Tozawa, Y. (2008). Identification and functional analysis of novel (p)ppGpp synthetase genes in Bacillus subtilis. Molecular microbiology 67, 291–304.

O’Farrell, P.H. (1978). The suppression of defective translation by ppGpp and its role in the stringent response. Cell 14, 545–557.

Ooga, T., Ohashi, Y., Kuramitsu, S., Koyama, Y., Tomita, M., Soga, T., and Masui, R. (2009). Degradation of ppGpp by nudix pyrophosphatase modulates the transition of growth phase in the bacterium Thermus thermophilus. J Biol Chem 284, 15549–15556.

Pausch, P., Steinchen, W., Wieland, M., Klaus, T., Freibert, S.A., Altegoer, F., Wilson, D.N., and Bange, G. (2018). Structural basis for (p)ppGpp-mediated inhibition of the GTPase RbgA. J Biol Chem 293, 19699–19709.

Pavlov, M.Y., Zorzet, A., Andersson, D.I., and Ehrenberg, M. (2011). Activation of initiation factor 2 by ligands and mutations for rapid docking of ribosomal subunits. EMBO J 30, 289–301.

Pereira, S.F., Gonzalez, R.L., Jr., and Dworkin, J. (2015). Protein synthesis during cellular quiescence is inhibited by phosphorylation of a translational elongation factor. Proc Natl Acad Sci U S A 112, E3274–3281.

Persky, N.S., Ferullo, D.J., Cooper, D.L., Moore, H.R., and Lovett, S.T. (2009). The ObgE/CgtA GTPase influences the stringent response to amino acid starvation in Escherichia coli. Molecular microbiology 73, 253–266.

Piir, K., Paier, A., Liiv, A., Tenson, T., and Maivali, U. (2011). Ribosome degradation in growing bacteria. EMBO Rep 12, 458–462.

Potrykus, K., Murphy, H., Philippe, N., and Cashel, M. (2011). ppGpp is the major source of growth rate control in E. coli. Environ Microbiol 13, 563–575.

Puszynska, A.M., and O’Shea, E.K. (2017). ppGpp Controls Global Gene Expression in Light and in Darkness in S. elongatus. Cell Rep 21, 3155–3165.

Remigi, P., Ferguson, G.C., McConnell, E., De Monte, S., Rogers, D.W., and Rainey, P.B. (2019). Ribosome Provisioning Activates a Bistable Switch Coupled to Fast Exit from Stationary Phase. Mol Biol Evol 36, 1056–1070.

Rittershaus, E.S., Baek, S.H., and Sassetti, C.M. (2013). The normalcy of dormancy: common themes in microbial quiescence. Cell host & microbe 13, 643–651.

Roelofs, K.G., Wang, J., Sintim, H.O., and Lee, V.T. (2011). Differential radial capillary action of ligand assay for high-throughput detection of protein-metabolite interactions. Proc Natl Acad Sci U S A 108, 15528–15533.

Rojas, A.M., Ehrenberg, M., Andersson, S.G., and Kurland, C.G. (1984). ppGpp inhibition of elongation factors Tu, G and Ts during polypeptide synthesis. Molecular & general genetics: MGG 197, 36–45.

Ross, W., Sanchez-Vazquez, P., Chen, A.Y., Lee, J.H., Burgos, H.L., and Gourse, R.L. (2016). ppGpp Binding to a Site at the RNAP-DksA Interface Accounts for Its Dramatic Effects on Transcription Initiation during the Stringent Response. Mol Cell 62, 811–823.

Schofield, W.B., Zimmermann-Kogadeeva, M., Zimmermann, M., Barry, N.A., and Goodman, A.L. (2018). The Stringent Response Determines the Ability of a Commensal Bacterium to Survive Starvation and to Persist in the Gut. Cell Host Microbe 24, 120–132 e126.

Shimizu, Y., Kuruma, Y., Kanamori, T., and Ueda, T. (2014). The PURE system for protein production. Methods in molecular biology 1118, 275–284.

Shyp, V., Tankov, S., Ermakov, A., Kudrin, P., English, B.P., Ehrenberg, M., Tenson, T., Elf, J., and Hauryliuk, V. (2012). Positive allosteric feedback regulation of the stringent response enzyme RelA by its product. EMBO Rep 13, 835–839.

Simonetti, A., Marzi, S., Myasnikov, A.G., Fabbretti, A., Yusupov, M., Gualerzi, C.O., and Klaholz, B.P. (2008). Structure of the 30S translation initiation complex. Nature 455, 416–420.

Srivatsan, A., Han, Y., Peng, J., Tehranchi, A.K., Gibbs, R., Wang, J.D., and Chen, R. (2008). High-precision, whole-genome sequencing of laboratory strains facilitates genetic studies. PLoS Genet 4, e1000139.

Stapels, D.A.C., Hill, P.W.S., Westermann, A.J., Fisher, R.A., Thurston, T.L., Saliba, A.E., Blommestein, I., Vogel, J., and Helaine, S. (2018). Salmonella persisters undermine host immune defenses during antibiotic treatment. Science 362, 1156–1160.

Steinchen, W., and Bange, G. (2016). The magic dance of the alarmones (p)ppGpp. Molecular microbiology 101, 531–544.

Steinchen, W., Vogt, M.S., Altegoer, F., Giammarinaro, P.I., Horvatek, P., Wolz, C., and Bange, G. (2018). Structural and mechanistic divergence of the small (p)ppGpp synthetases RelP and RelQ. Sci Rep 8, 2195.

Svitil, A.L., Cashel, M., and Zyskind, J.W. (1993). Guanosine tetraphosphate inhibits protein synthesis in vivo. A possible protective mechanism for starvation stress in Escherichia coli. J Biol Chem 268, 2307–2311.

Szaflarski, W., and Nierhaus, K.H. (2007). Question 7: optimized energy consumption for protein synthesis. Orig Life Evol Biosph 37, 423–428.

Tagami, K., Nanamiya, H., Kazo, Y., Maehashi, M., Suzuki, S., Namba, E., Hoshiya, M., Hanai, R., Tozawa, Y., Morimoto, T., et al. (2012). Expression of a small (p)ppGpp synthetase, YwaC, in the (p)ppGpp(0) mutant of Bacillus subtilis triggers YvyD-dependent dimerization of ribosome. Microbiologyopen 1, 115–134.

Tempest, D.W., and Neijssel, O.M. (1984). The status of YATP and maintenance energy as biologically interpretable phenomena. Annu Rev Microbiol 38, 459–486.

Verstraeten, N., Fauvart, M., Versees, W., and Michiels, J. (2011). The universally conserved prokaryotic GTPases. Microbiology and molecular biology reviews: MMBR 75, 507–542.

Vinogradova, D.S., Kasatsky, P., Maksimova, E., Zegarra, V., Paleskava, A., Konevega, A.L., and Milon, P. (2019). How the initiating ribosome copes with (p)ppGpp to translate mRNAs. bioRxiv, 545970.

Wagner, E.G., and Kurland, C.G. (1980). Translational accuracy enhanced in vitro by (p)ppGpp. Molecular & general genetics: MGG 180, 139–145.

Wang, B., Dai, P., Ding, D., Del Rosario, A., Grant, R.A., Pentelute, B.L., and Laub, M.T. (2019). Affinity-based capture and identification of protein effectors of the growth regulator ppGpp. Nat Chem Biol 15, 141–150.

Wang, J., Caban, K., and Gonzalez, R.L., Jr. (2015). Ribosomal initiation complex-driven changes in the stability and dynamics of initiation factor 2 regulate the fidelity of translation initiation. Journal of molecular biology 427, 1819–1834.

Wang, J.D., Sanders, G.M., and Grossman, A.D. (2007). Nutritional control of elongation of DNA replication by (p)ppGpp. Cell 128, 865–875.

Wienk, H., Tishchenko, E., Belardinelli, R., Tomaselli, S., Dongre, R., Spurio, R., Folkers, G.E., Gualerzi, C.O., and Boelens, R. (2012). Structural dynamics of bacterial translation initiation factor IF2. J Biol Chem 287, 10922–10932.

Zhang, Y., Zbornikova, E., Rejman, D., and Gerdes, K. (2018). Novel (p)ppGpp Binding and Metabolizing Proteins of Escherichia coli. MBio 9.

